# Dual targeting of MAPK and PI3K pathways unlocks redifferentiation of *Braf*-mutated thyroid cancer organoids

**DOI:** 10.1101/2023.03.30.534915

**Authors:** Hélène Lasolle, Andrea Schiavo, Adrien Tourneur, Pierre Gillotay, Bárbara de Faria da Fonseca, Lucieli Ceolin, Olivier Monestier, Benilda Aganahi, Laura Chomette, Marina Malta Letro Kizys, Lieven Haenebalcke, Tim Pieters, Steven Goossens, Jody Haigh, Vincent Detours, Ana Luiza Silva Maia, Sabine Costagliola, Mírian Romitti

## Abstract

Thyroid cancer is the most common endocrine malignancy and several genetic events have been described to promote the development of thyroid carcinogenesis. Besides the effects of specific mutations on thyroid cancer development, the molecular mechanisms controlling tumorigenesis, tumor behavior, and drug resistance are still largely unknown. Cancer organoids have been proposed as a powerful tool to study aspects related to tumor development and progression and appear promising to test individual responses to therapies. Here, using mESC-derived thyroid organoids, we developed a Braf^V637E^- inducible model able to recapitulate the features of papillary thyroid cancer *in vitro*. Overexpression of the murine Braf^V637E^ mutation, equivalent to Braf^V600E^ in humans, rapidly triggers to MAPK activation, cell dedifferentiation, and disruption of follicular organization. Braf^V637E^-expressing organoids show a transcriptomic signature for p53, focal adhesion, ECM-receptor interactions, EMT, and inflammatory signaling pathways. Finally, PTC-like thyroid organoids were used for drug screening assays. The combination of MAPK and PI3K inhibitors reversed *Braf*^V637E^ oncogene-promoted cell dedifferentiation while restoring thyroid follicle organization and function *in vitro*. Our results demonstrate that pluripotent stem cells-derived thyroid cancer organoids can mimic tumor development and features while providing an efficient tool for testing novel targeted therapies.

## INTRODUCTION

Pluripotent Stem Cells (PSCs) emerged as a model to dissect and recapitulate the molecular events and gene networks that regulate cell fate determination, cell differentiation and organogenesis. Guided differentiation of PSCs into 3D organized tissue by inducible overexpression of specific transcription or growth factors and inhibitors allows the derivation of *in vitro* organoids and the understanding of mechanisms regulating cell differentiation and tissue formation^1^. Due to their remarkable self-organizing structures and functional properties, organoid technology became a powerful tool to model organ development and disease ‘in a dish’ ^2,3^.

Our group has demonstrated the generation of functional thyroid organoids derived from mouse and human embryonic stem cells (mESC and hESC)^4,5^. The differentiation protocols rely on transient induction of *Nkx2.1* and *Pax8* thyroid transcription factors. This approach enables an efficient generation of thyroid follicular cells that organize into three-dimensional follicular structures capable of thyroid hormone production *in vitro* and *in vivo*. These models, along with the further development of thyroid organoids derived from healthy murine and human pluripotent or adult stem cells, (PSCs/aSC)^6–16^ open new perspectives to explore thyroid developmental and pathological processes, including thyroid cancer.

In the last decade, the use of organoids for cancer research emerged opening new possibilities to better understand tumor behavior. Initially, colon, prostate, pancreatic, ovarian, lung and thyroid cancer-derived organoids have been efficiently generated, which resemble phenotypically and genetically the tumor of origin^17–26^. A large set of healthy and tumour-matching organoids has been generated and is available through biobanks. These patient-derived organoids are particularly interesting in testing individual responses to therapies^21,26,27^.

Recently, studies have reported the generation of cancer models arising from healthy (aSCs and PSCs) cells by controlling oncogene expression^28–30^. By using shRNA and CRISPR-Cas9, stomach cancer (*Cdh1^-/-^; Tp53^-/-^)*^31^, colon cancer (*APC, TP53, KRAS* and *Smad4* mutations)^28,29,32^, pancreatic cancer (*KRAS* and/or *TP53* mutations)^17,33^ and lung adenocarcinoma organoids (*HER2* overexpression) could be efficiently generated^30^. Compared to patient-derived organoids, healthy stem cells-derived cancer organoids are suitable to address additional questions such as precise effects of oncogenes and early events driving tumorigenesis; the role of cancer stem cells on tumor induction, genomic stability, the effect of treatments at different stages of carcinogenesis and screening for new therapeutical tools^34^. Besides, they allow multiple reproducible experiments free from inter-individual variability.

Thyroid cancer is the most frequent endocrine malignancy, and several genetic events have been described as driving thyroid carcinogenesis. Papillary thyroid carcinoma (PTC) is the most common malignant thyroid tumor^35^, and aberrant activation of the MAPK pathway is a hallmark^36^. The BRAF^V600E^ mutation is the most frequent genetic event, accounting for around 50% of PTCs^37–39^ and results in an oncogenic protein with markedly elevated kinase activity that constitutively induces MEK/ERK signaling transduction^40–42^.

*BRAF*-mutated thyroid tumors are often less differentiated, mainly due to the lower expression levels of thyroid functional genes *NIS, TSHR, TG* or *TPO,* leading to a low Thyroid Differentiation Score (TDS)^43^. This dedifferentiated state is associated with a worse clinical condition and with a higher rate of radioactive iodine (RAI) refractory tumors^44,45^. However, its impact on prognosis is still uncertain, as BRAF^V600E^ alone has not been proven to be an independent prognostic factor^46,47^. On the other hand, *BRAF* and *TERT* promoter mutated tumors are associated with higher specific mortality^48^. Also, studies have shown a small proportion (around 20%) of BRAF^V600E^ mutated tumors presenting a higher level of differentiation associated with less aggressive behavior and a preserved RAI uptake ability^43,49^. This suggests a possible cell heterogeneity among the *BRAF^V^*^600^*^E^*-mutated tumors.

So far, efficient, fast, and cost-effective tools are lacking to functionally characterize a large set of candidate genes in the context of thyroid carcinogenesis. Thyroid cancer organoids/spheroids models have been recently described based on patient tumor-derived models^50–54^. Tumor-derived organoids can be used to explore patient-specific tumor behavior and the treatment response, but present high heterogeneity and low reproducibility. Saito et al. described a mouse model of poorly differentiated thyroid carcinoma obtained after transplantation of thyroid organoids derived from *Tp53*^-/-^ mice with *NRAS* activating mutation^25^. Recently, Veschi *et al.* ^55^ generated PTC and FTC organoids by inducing *BRAF*, *NRAS* and *TP53* mutations in thyroid progenitors derived from hESC. After transplantation, the tumors resembled Papillary and Follicular tumors and, transcriptomics analysis suggested a cooperative effect of Kisspeptin receptor (KISS1R) and Tissue Inhibitor of Metalloproteinase 1 (TIMP1)/Matrix metallopeptidase 9 (MMP9)/Cluster of differentiation 44 (CD44), on tumor development and progression^55^.

In this study, we have established an *in vitro* organoid model for PTC to streamline the process of screening potential targets and pharmaceutical agents. We achieved this by leveraging our established model of thyroid generation, which involves the inducible expression of the Braf^V637E^ mutation in functional thyroid follicles derived from mouse ESCs.

## RESULTS

### mESC_BRAF^V637E^ cell line generation and characterization

We combined the previously described mESC transient induction thyroid differentiation strategy^4^ with a Braf^V637E^-ERT^2^ inducible model^56^ to generate a double-inducible recombinant mESC_Braf^V637E^ cell line. The murine Braf^V637E^ mutation corresponds to the human Braf^V600E^ mutation. The resulting mESC_Nkx2-1/Pax8_bTg_Braf^V637E^_ERT^2^ clones were initially selected and characterized by the response to doxycycline (Dox) treatment, confirmed by the overexpression of Nkx2-1 and Pax8; and ability to spontaneously differentiate into cells from the three germ-layers (Supplementary Fig. 1A-C). In addition, a control mESC line was generated in which Braf^V637E^_ERT^2^ system was replaced by the eGFP sequence (mESC_Nkx2-1/Pax8_bTg_eGFP). Similarly, the control line responded properly to Dox induction and maintained the pluripotency capacity (Supplementary Fig. 1D-F).

### Thyroid differentiation protocol

The capability of the new cell line to generate thyroid follicles and express the Braf^V637E^ oncogene was demonstrated following the mESC thyroid differentiation protocol^4,57,58^. Since we used a newly modified mESC line, the steps of the differentiation protocol were tested and adapted accordingly (Fig. 1A). Here, using the mESC_Nkx2-1/Pax8_bTg_Braf^V637E^ line, we observed that Dox stimulation for 5 days leads to higher exogenous and endogenous *Nkx2-1*, *Pax8* and *Tg* mRNA compared to 3 or 4 days of treatment (Supplementary Fig. 2A-D). Similar results were obtained using the control mESC line (data not shown). Next, two weeks of treatment with cAMP (from day 9 to 23) markedly induced the expression levels of *Nkx2-1, Pax8, Tg, TshR, Nis, Tpo* and exogenous *Braf^V^*^637^*^E^* when compared to the control (-Dox -cAMP) (Fig. 1B). Immunofluorescence staining also showed Nkx2-1 and Tg-expressing cells (Fig. 1C) organized into follicular structures with intraluminal iodinated-Tg (Tg-I) accumulation (Fig. 1D). Since our protocol also generates other cell types than thyroid^58^, a follicle enrichment (FE) step was added to the protocol to enrich the follicular structures and induce the Braf^V637E^ oncogene in a purer population. This protocol led to an increase in thyroid markers levels (Fig. 1E) when compared to the non-enriched control condition (No FE), while the follicular structures and functionality were preserved as demonstrated by Nkx2-1, Nis and Tg-I stainings (Fig. 1F-G). Notably, after re-embedding in Matrigel (MTG), follicles could be maintained in culture for at least 28 additional days without major histological changes (data not shown).

**Fig. 1.**
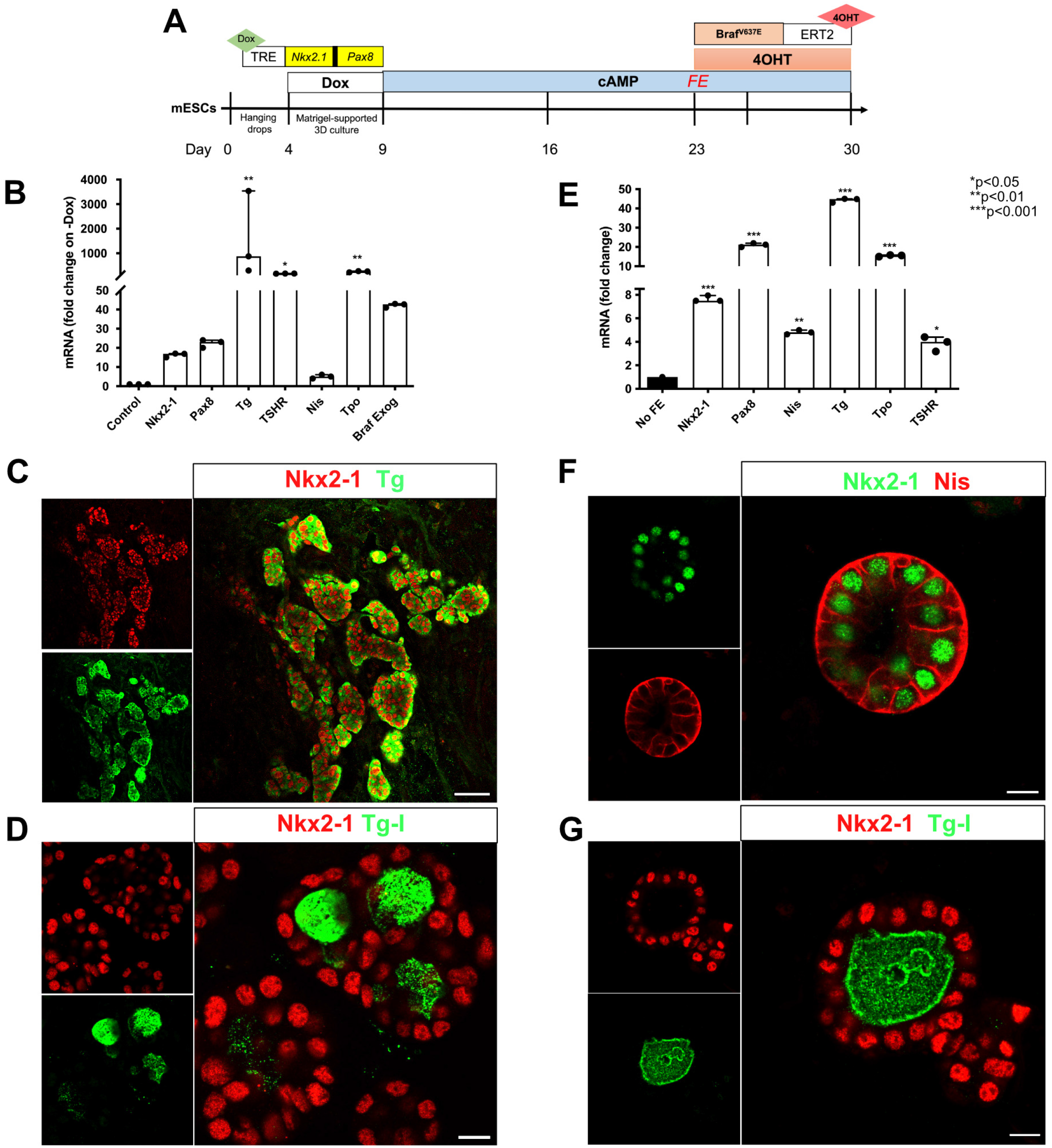
mESC_Nkx2-1/Pax8_bTg_Braf^V637E^ cell line differentiation into thyroid organoids. Scheme of the thyroid differentiation protocol and Braf^V637E^ induction in mature thyroid follicles (A). Differentiation of mESC_Nkx2-1/Pax8_bTg_Braf^V637E^ cell line promotes expression of the main thyroid genes and Braf^V637E^ exogenous (B). Control corresponds to the - Dox condition. Immunofluorescence staining showing Nkx2-1 and Tg co-expressing cells organized in follicular structures (C), which are accumulating the thyroid hormone precursor, Tg-I, inside the lumen compartment (D). The follicular enrichment (FE) procedure significantly increased the expression levels of thyroid genes (E) while keeping the structural organization of the follicles evidenced by Nis basolateral localization (F) and its functionality, with Tg-I accumulation (G). Values represent the median (IQR) of 3 independent experiments with individual values shown (*p<0.05 ; **p<0.01 ; ***p<0.001; Mann-Whitney U test). Scale bars, 50 μm and 10 μm for high magnification follicles.

### Braf^V637E^ induction on thyroid organoids leads to PTC-like phenotype

To ensure that Braf^V637E^ oncogene is expressed specifically among follicular thyroid cells, we used the bovine Tg (bTg) as a promoter controlling Braf^V637E^_ERT^2^ expression. Effectively, exogenous *Braf^V^*^637^*^E^*mRNA expression was rapidly induced after Dox treatment (Day 9; Supplementary Fig. 2E), maintaining the expression levels stable over time (Fig.1B and Supplementary Fig. 3A), while *Tg* expression increased following the differentiation program (Supplementary Fig. 3A). In the absence of tamoxifen (4OHT) Braf^V637E^ protein is not active/phosphorylated since it is maintained in the nucleus in an inactive complex with HSP90. The addition of 4OHT induces nuclear translocation of the Braf^V637E^ mutant to the cytoplasm (NES signal) ^56^, then driving MAPK activation^56^ (Fig. 2A). Indeed, 48 hours after 4OHT addition, we observed an increase in ERK phosphorylation (pERK) compared to the control condition (cAMP) (Fig. 2B).

**Fig. 2.**
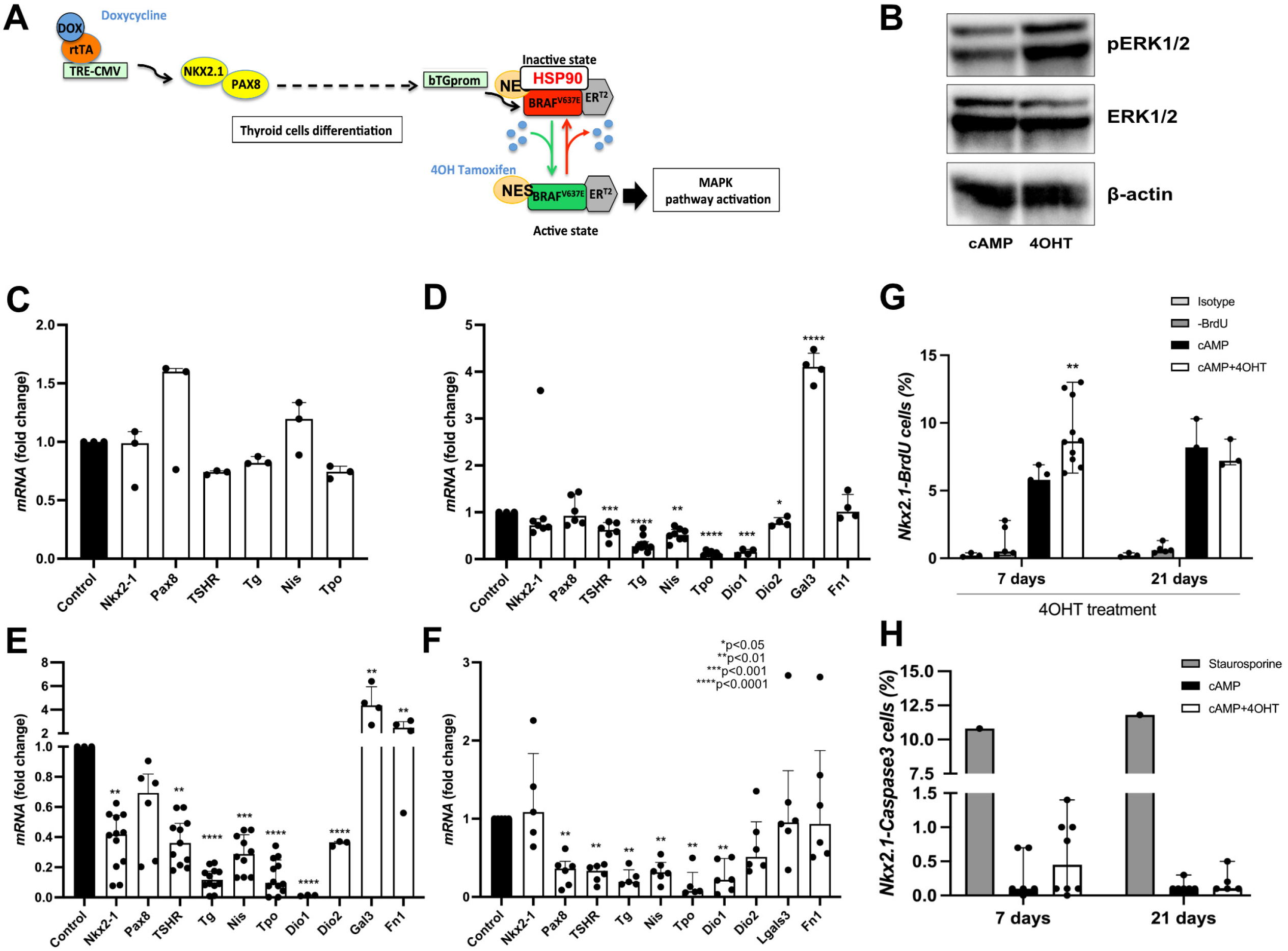
Effect of the Braf^V637E^ oncogene induction on mature thyroid follicles. Schematic representation of thyroid differentiation, Braf^V637E^ oncogene expression on thyroid cells, and its activation under tamoxifen (4OHT) treatment (A). Western blot demonstrates an increase of phospho-ERK (pERK) 48 hours after the addition of 4OHT to the organoids culture (B). Beta-actin was used as a loading control for the immunoblot experiments. The image represents one experiment from 3 experimental replicates. Gene expression analysis showing the inhibitory effect of Braf^V637E^-oncogene activation on thyroid genes after 6 hours (C), 48 hours (D), 7 days (E) and 21 days (F) of 4OHT treatment. For each time point gene expression levels of the cAMP+4OHT treated cells were compared to the control (cAMP) levels. Bar graphs represent the median (IQR) of at least 3 independent experiments with individual values shown. (*p<0.05 ; **p<0.01 ; ***p<0.001; ***p<0.0001; Mann-Whitney U test). Proportions of proliferating (Nkx2.1/BrDU+) (G) and apoptotic (Nkx2.1/Caspase3+) (H) cells among control (cAMP) and Braf^V637E-^induced (4OHT) organoids after 7 and 21 days of 4OHT treatment. For the proliferation assay, isotype and -BrdU conditions were used as negative controls for flow cytometry gating. As a positive control for apoptosis, staurosporine treatment (24 h) was used to determine Caspase 3 expression. Bar graphs represent the median (IQR) of at least 3 independent experiments with individual values shown. cAMP was used as control for comparisons. (**p<0.01; Mann-Whitney U test).

One of the hallmarks of thyroid cancer is the downregulation of thyroid differentiation markers^41,59^. In our organoid model, the activation of Braf^V637E^ rapidly led to a progressive time-dependent decrease of *Tpo*, *TSHR* and *Tg* mRNA expression (Fig. 2C-E and Supplementary Fig. 3B), detected as early as 6 hours after 4OHT addition (Fig. 2C). Notably, more significant downregulation compared to the cAMP control was observed after 48 hours and 7 days while maintained for at least 21 days (Fig. 2D-F, respectively). As for PTCs, in our model, the expression of thyroid transcription factors *Pax8* and *Nkx2-1* was also globally maintained. However, a partial decrease of *Nkx2.1* levels was observed at 7 days, which corroborates recent findings suggesting that thyroid cells transiently downregulated NKX2.1 in early tumor stages^60^. Conversely, Galectin 3 *(Lgals3)* expression, which has been suggested as a marker for thyroid malignancies and, more specifically, for PTC^61^, was significantly increased at 48 h and 7 days of 4OHT treatment (Fig. 2D-E). Also, an increase in Fibronectin 1 (*Fn1*) expression, associated with aggressive thyroid cancer^62,63^, was observed after 7 days of Braf^V637E^ induction, suggesting a more advanced dedifferentiated cell state in our organoid model (Fig. 2E). The generation of thyroid organoids was equally successful when differentiating mESC_Nkx2-1/Pax8_bTg_Braf^V637E^_ERT2 cells using hrTSH instead of cAMP, as demonstrated in Supplementary Figure 4A and C. Furthermore, the addition of 4OHT to the conditioned media (for a duration of 7 days, as illustrated in Supplementary Figure 4B-C) led to dedifferentiation and histological changes.

Cell proliferation assessment demonstrated an increased proportion of Nkx2-1/BrdU+ cells within the Braf^V637E^-expressing organoids (4OHT) as compared to the controls (cAMP) after 7 days of treatment (Fig. 2G). Interestingly, there were no observable variations in proliferation even after 21 days of 4OHT treatment (Fig. 2G). Furthermore, considering that prior studies have indicated that the induction of oncogenes in organoids can lead to cell death^64^, our analysis of apoptosis showed no significant difference in the proportions of Nkx2.1/Caspase3+ between cAMP and 4OHT-treated organoids both at 7 and 21 days of 4OHT treatment (Fig. 2H). However, a notable increase in the proportion of Nkx2.1/Caspase3+ cells was evident when cAMP-treated organoids were exposed for 24h to staurosporine, a highly potent inducer of apoptosis. (Fig. 2H). Given that tumor growth arrest and antiapoptotic phenotype are linked to oncogene-induced senescence (OIS)^65,66^, we examined p21 expression at both 7 and 21 days of 4OHT treatment. Notably, we observed no co-expression of p21 with Nkx2-1 in either the control or 4OHT-treated thyroid organoids. However, a small proportion of non-thyroid cells (Nkx2.1 negative) were positive for p21 and used as positive controls (Supplementary Fig. 5A-C).

Histological characterization performed after 48 hours of continuous activation of Braf^V637E^ by 4OHT demonstrated disruption of the follicular organization in non-enriched thyroid organoids compared to the control condition (cAMP) (Fig. 3A). Nkx2-1 and Tg staining revealed that Braf^V637E^ oncogene strongly disturbs thyroid follicles resulting in elongated and unorganized structures. In addition, heterogeneous expression of Tg was observed among the Nkx2-1 cells (Fig. 3A). Further analysis, performed in follicle enriched (FE) population, confirmed the effect of Braf^V637E^ oncogene in promoting cell dedifferentiation and loss of follicle organization. Tg levels were overall reduced and heterogeneous among the unorganized Nkx2-1+ cells (Fig. 3B). Interestingly, the cell disorganization caused by Braf^V637E^ oncogene might pass by an initial expansion of the follicular size with infiltration of Nkx2-1 cells inside of the lumen compartment, which also presents a heterogeneous distribution of Tg (Supplementary Fig. 3C) since this feature was observed among some “follicular-like” remaining structures after 48 h of 4OHT treatment. Reassuringly, staining for E-Cadherin (epithelial) and Zo-1 (intra-luminal) adhesion markers showed that Braf^V637E^-expressing cells are not able to preserve polarization and, consequently lumen space is not well defined in most of the structures (Fig. 3C and D, respectively). Therefore, the functionality of those cells was impaired and, Nis expression and Tg-I accumulation were significantly reduced compared to the control condition (Fig. 3E-F). Of note, despite the strong effect of Braf^V637E^ oncogene on thyroid follicle disruption, we observed that few follicles still preserved a certain degree of organization (Supplementary Fig. 3D) after 48 h of 4OHT treatment. As expected, after 7 days of Braf^V637E^ continuous stimulation, a higher level of cell dedifferentiation and lack of organization could be detected, with a lower proportion of the Nkx2-1 cells expressing Tg compared to control (cAMP) (Fig. 3G) and to the previous time point (48 h; Fig. 3A). However, despite the absence of proper follicular organization, there was an increased proportion of Tg-expressing cells within the Braf^V637E^-expressing organoids after 21 days of 4OHT treatment compared to earlier time points (48 h and 7 days; Fig. 3H). This observation, in conjunction with the gene expression and the proliferation data (Fig. 2F-G) implies a partial arrest in the tumorigenic processes at a later stage.

**Fig. 3.**
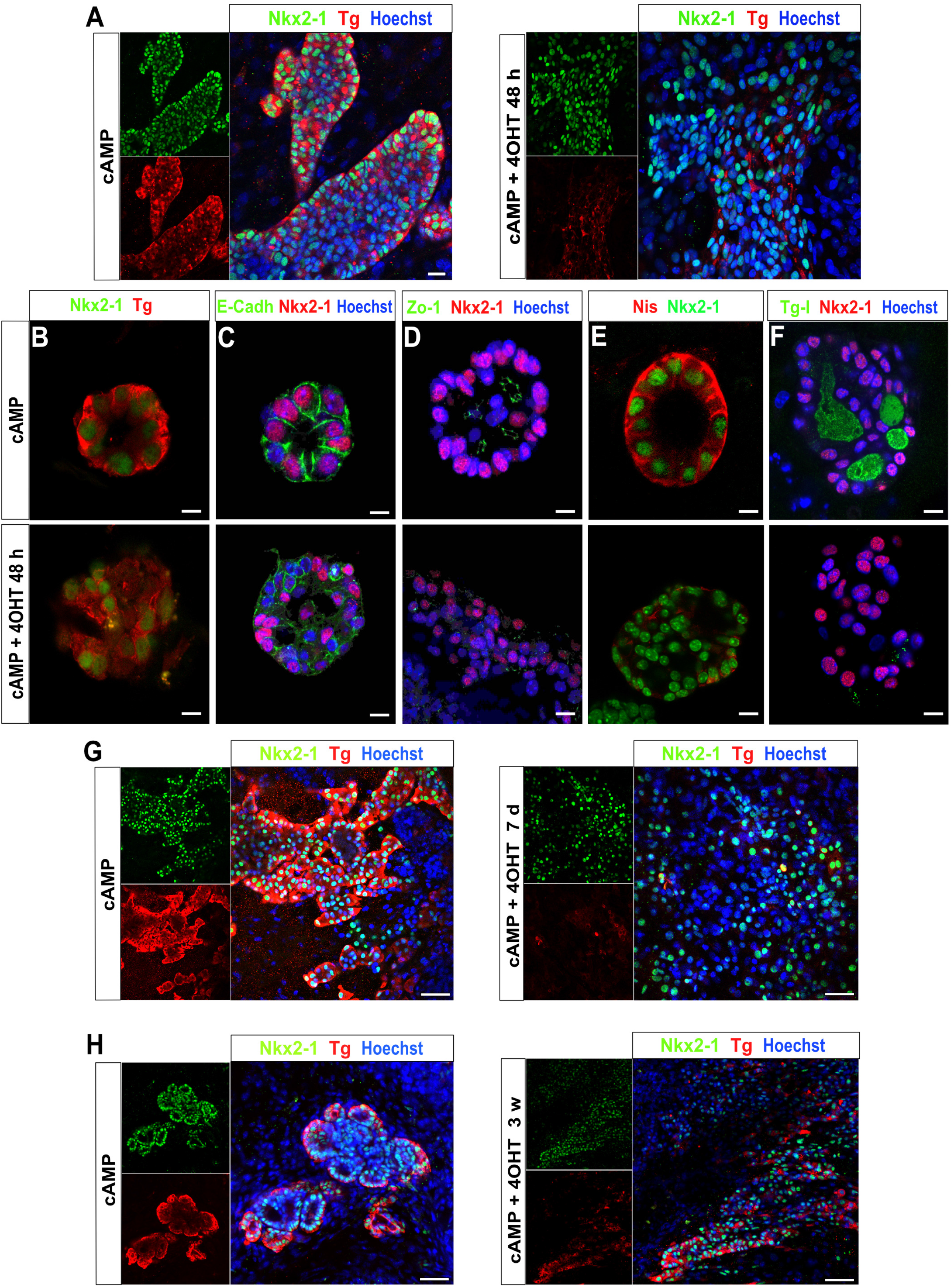
Morphological changes on thyroid follicles caused by Braf^V637E^ activation. cAMP-treated thyroid cells show follicular organization with Nkx2-1 nuclear expression and Tg accumulation in the luminal compartment. In contrast, after 48 h of Braf^V637E^ oncogene induction by 4OHT, most of the cells are not organized into follicular structures and a great proportion is expressing very low levels of Tg (A). Higher magnification images showing the follicular organization of the thyroid cells in the control condition (48 h), with proper expression of Nkx2-1 and/or Tg (B), E-cadherin (C), Zo-1 (D), Nis (E) and Tg-I (F) accumulation in the lumen. While in the 4OHT condition the follicular organization is disrupted as well as its function. Nkx2-1 and Tg co-staining in control (cAMP) and Braf^V637E^-induced (4OHT) cells for 7 days (G) and 21 days (H) shows clear changes in thyroid morphology evidenced by the heterogeneity of Nkx2-1 cells which mostly do not express Tg or at low levels at day 7, while at day 21 a higher proportion of Nkx2-1 cells are Tg positive. Hoescht (shown in blue) was used for nuclei staining. Scale bars, 20 μm (A), 10 μm (B-F) and 50 μm (G-H).

The above-described experiments were also performed using the control TRE-Nkx2-1-Pax8_bTg-eGFP cell line. Adding 4OHT to mature follicles for 7 days did not impair the levels of the thyroid differentiation markers and the follicles’ organization (Supplementary Fig. 6A-B). Furthermore, there was a slight increase in the proportions of Nkx2.1 and GFP-expressing cells when WT organoids were treated with 4OHT (7 days; Supplementary Fig. 6C). It’s worth noting, as previously demonstrated^58^, that bTg_GFP-expressing cells do not encompass the entire Tg-expressing cells population (Supplementary Fig. 6D). This could be attributed to the lack of regulatory regions not included in our construct.

### Effect of transient Braf^V637E^-induction on cell differentiation

Since our system for Braf^V637E^ activation is dependent on continuous treatment with 4OHT, we transiently treated the cells with 4OHT for 2 days, followed by 5 days of culture in a 4OHT-free medium (cAMP) in order to explore if the oncogenic effect is maintained when the protein is not active. Surprisingly, we did not observe recovery in *Nkx2-1, TSHR, Tg* or *Tpo* gene expression levels (Supplementary Fig. 7A, C-E) after 4OHT removal. However, *Pax8* and *Slc5a5/Nis* expression was partially recovered (Supplementary Fig. 7B and F), confirming the specific inhibitory effect of Braf^V637E^ oncogene on *Slc5a5/Nis* expression. These findings indicate that *Nis* regulatory mechanisms are maintained in our system and suggest it as a tool for Nis-reactivation studies.

### Screening for Nis re-expression using signaling pathways inhibitors

Studies in mouse models and humans have demonstrated that tumors carrying *BRAF* mutations show reduced expression of *Nis* and, consequently a higher rate of radioiodine (RAI) refractoriness^41,44,67^. Here, we used our organoids model to explore new strategies to reactivate *Nis* expression in Braf^V637E^-expressing cells by screening distinct categories of inhibitors previously described to be involved in *Nis* regulation ^68,69^. PD0325901 (MEK inhibitor), LY204002 (PI3K inhibitor), VPA (HDAC inhibitor), NAC (N-acetyl cysteine, antioxidant compound), 5-AZA-2’ deoxycytidine and RG108 (DMNT inhibitors) were tested isolated and/or in combination for 3 days (in addition to cAMP+4OHT) using Braf^V637E^-expressing organoids (previously treated for 4 days with 4OHT; Fig. 4A). Initially, we observed that VPA alone was the only treatment able to completely restore *Nis* expression levels (Fig. 4B, C and Supplementary Fig. 8A). However, when combined, PD0325901 and LY204002 inhibitors; PD0325901, LY204002 and VPA; and PD0325901 and NAC resulted in a great increase of *Nis* mRNA (Fig. 4B, C and Supplementary Fig. 8A). IF staining shows Nis protein re-expression among Braf^V637E^-induced cells treated with VPA alone, but localization was not restricted to the basolaterall membrane. In contrast, when cells were treated with MEK (PD) of PI3K (LY) inhibitors, associated or not with VPA, we observed Nis protein correctly localized at the basolateral membrane and surprisingly, it restored the follicular structure (Fig. 4C).

**Fig. 4.**
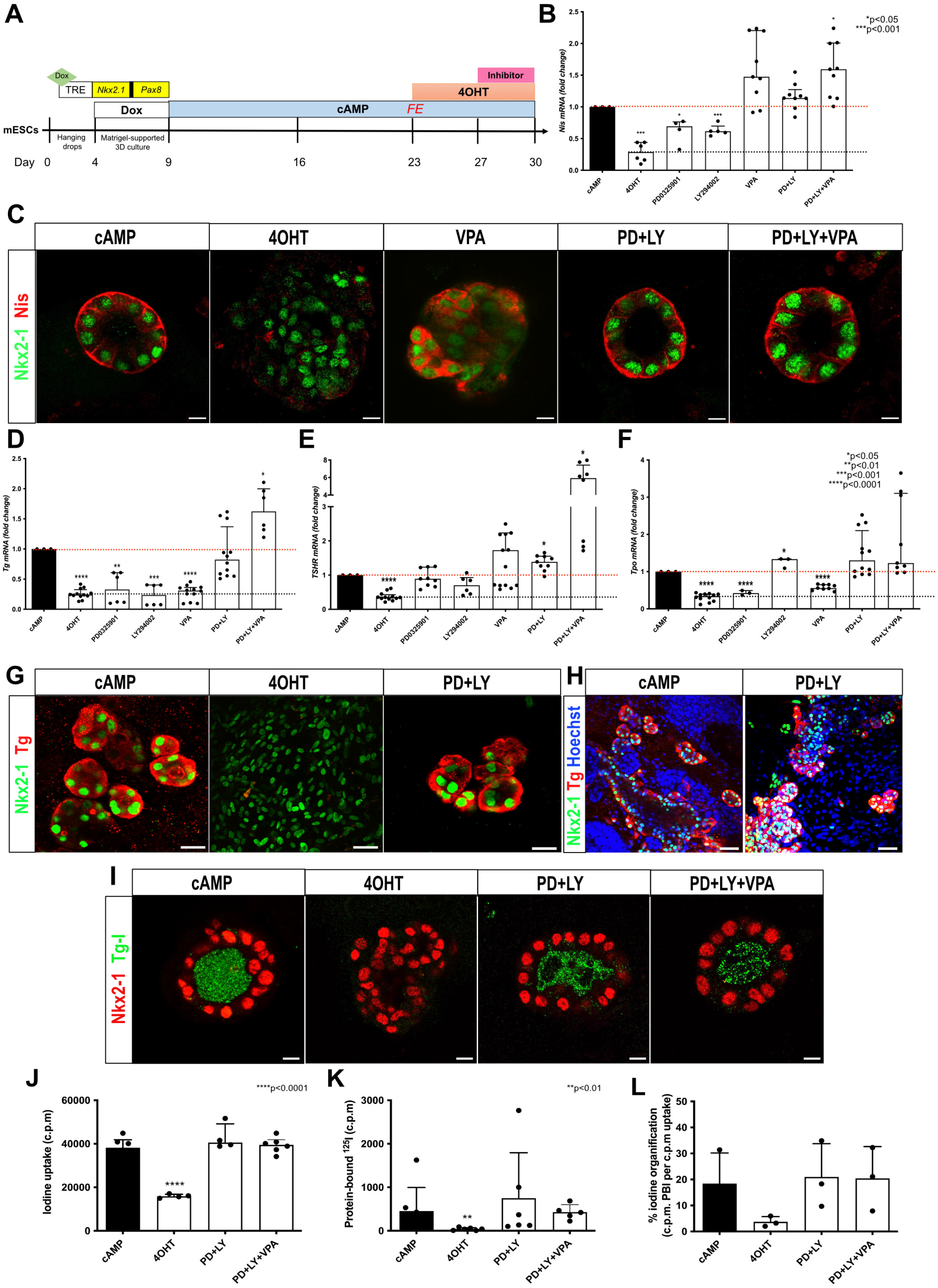
Drug screening reveals that VPA and MAPK/PI3K inhibition can restore Nis expression in Braf^V637E^-induced cells. Schematic representation of the protocol used for drug screening experiments (A). qPCR data show that 4OHT-treated organoids (4 days) treated with MEK (PD0325901; PD) and PI3K (LY294002; LY) inhibitors isolated increase but do not restore *Nis* expression to control (cAMP) levels. However, when combined, *Nis* expression reached cAMP levels. On the other hand, the HDAC inhibitor (VPA) can recover *Nis* expression (B) by itself. Bar graphs represent the median (IQR) of at least 4 independent experiments with individual values shown. Comparisons were performed against the cAMP condition. (**p<0.05 ; ***p<0.001 ; Mann-Whitney U test). Confocal images show downregulation of the Nis transporter in 4OHT condition, which is restored by VPA treatment. However, the expression pattern differs from the control (cAMP) condition. Co-inhibition of MAPK and PI3K pathways associated or not to VPA treatment restores Nis protein expression at the basolateral membrane of the properly organized follicles (C). Scale bars, 10 μm. Gene expression analysis shows that PD0325901 and LY294002 co-treatment recover the *Tg* (D), *TSHR* (E), and *Tpo* (F) mRNA to cAMP levels. Bar graphs represent the median (IQR) of at least 3 independent experiments with individual values shown. Comparisons were performed against the cAMP condition. (*p<0.05 ; **p<0.01 ; ***p<0.001; ***p<0.0001; Mann-Whitney U test). Immunostaining for Nkx2-1 and Tg shows that proteins levels are similar to controls (cAMP) while follicular organization (G, H) and luminal Tg-I accumulation were restored under PD+LY and PD+LY+VPA conditions (I). Scale bars, 20 μm (G), 50 μm (H), and 10 μm (I). Organification assay shows recovery of Iodine uptake (J), Protein-bound to 125I (K), and % of iodine Organification (L) in PD+LY and PD+LY+VPA conditions. Bar graphs represent the median (IQR) of at least 3 independent experiments with individual values shown. Comparisons were performed against the cAMP condition. (**p<0.01 ; ***p<0.0001; Mann-Whitney U test).

### MAPK and PI3K signaling inhibition cause Braf^V637E^-induced cell redifferentiation and function recovery

Since we observed that HDAC, MEK, and PI3K inhibition seem to favor follicle reorganization, we evaluated if such treatments also induce cell redifferentiation and function recovery. Initially, we analyzed the gene expression levels of thyroid differentiation markers strongly affected by Braf^V637E^ induction. The effects of isolated and combined drugs on *Tg, TSHR, Tpo, Nkx2-1* and *Duox* genes appear to be variable and pathway-dependent (Supplementary Fig. 8B-F). As for *Slc5a5/Nis*, MEK and PI3K inhibition in combination with or without VPA resulted in complete recovery of *Tg*, *TSHR* and *Tpo* mRNA levels (Fig. 4D-F), indicating cell redifferentiation. While *TSHR* levels can be restored by treatment with isolated inhibitors (Fig. 4E), *Tpo* expression appears to be dependent on PI3K signaling (Fig. 4F). On the other hand, *Tg* mRNA expression was recovered only when the inhibitors were used in combination (Fig. 4D). Indeed, immunostaining showed that isolated inhibitors could not restore Tg levels. Nevertheless, a higher proportion of Tg-expressing cells was observed with the MEK inhibitor PD0325901 (Supplementary Fig. 8G). Conversely, combined MEK and PI3K inhibitors restored Tg protein levels and led to a significant reorganization of the cells into follicles comparable to the control (cAMP) condition (Fig. 4G-H). Considering that inhibition of MEK and PI3K pathways (+/-VPA) recovered thyroid differentiation and follicular organization, we tested the functionality of reorganized follicles. Remarkably, we observed an accumulation of Tg-I in the lumen of follicles treated with the combinations of inhibitors (Fig. 4I). The organification assay confirmed that the co-treatment restored iodide uptake and ^125^I binding to proteins resulting in organification levels comparable to the control (Fig. 4J-L).

### Dabrafenib and trametinib effect on redifferentiation is potentialized by PI3K inhibition

The treatment of advanced thyroid carcinomas as PDTC and ATC with BRAF inhibitor dabrafenib and MEK inhibitor trametinib have shown significant redifferentiation and response rates in *BRAF*-mutated tumors^70–72^. In this study, we examined the effect of isolated dabrafenib and trametinib inhibitors, as well as their combination with PI3K inhibition, on thyroid cancer organoids. Using the same experimental approach (Fig. 4A), we noted that both dabrafenib and trametinib treatments successfully restored *Nis* expression to levels comparable to the control group (treated with cAMP). Notably, the combination of trametinib with a PI3K inhibitor yielded an even more significant recovery, resulting in a threefold increase compared to the control group (Fig. 5A). Furthermore, isolated treatments with dabrafenib and trametinib elicited a moderate restoration of the main thyroid differentiation markers’ expression. However, the most substantial recovery of *Tg, Tpo*, and *TSHR* levels was observed when organoids were subjected to a combination of dabrafenib and trametinib along with a PI3K inhibitor, with the latter showing the most pronounced effect (Fig. 5B-D). Interestingly, both Tg and Nis proteins were detectable in all experimental conditions, with some cells organized into functional follicles able to produce Tg-I (Fig. 5E). However, due to the complexity of the cell composition and organization of our 3D organoids, accurately quantifying the proportion of reorganized follicles proved to be very challenging and could not be efficiently assessed in the present study. Furthermore, our proliferation and apoptosis assays revealed that co-treatment with Trametinib and LY led to a decrease in proliferation (Nkx2.1/BrdU+ cells) and a marginal increase in apoptosis (Nkx2.1/Caspase 3 + cells). Comparable outcomes were achieved through incubation with MEK and PI3K inhibitors (PD+LY). Nevertheless, it’s worth noting that apoptosis induction was notably more pronounced when compared to the effects observed with Trametinib+LY treatment (Fig. 5F-G). Western blot analysis distinctly revealed an increase in pERK levels after a 7-day 4OHT treatment. However, when Trametinib and LY were combined in the treatment, it led to a mild inhibition of pERK compared to the conditions involving 4OHT alone or in combination with PD+LY (Fig. 5H).

**Fig. 5.**
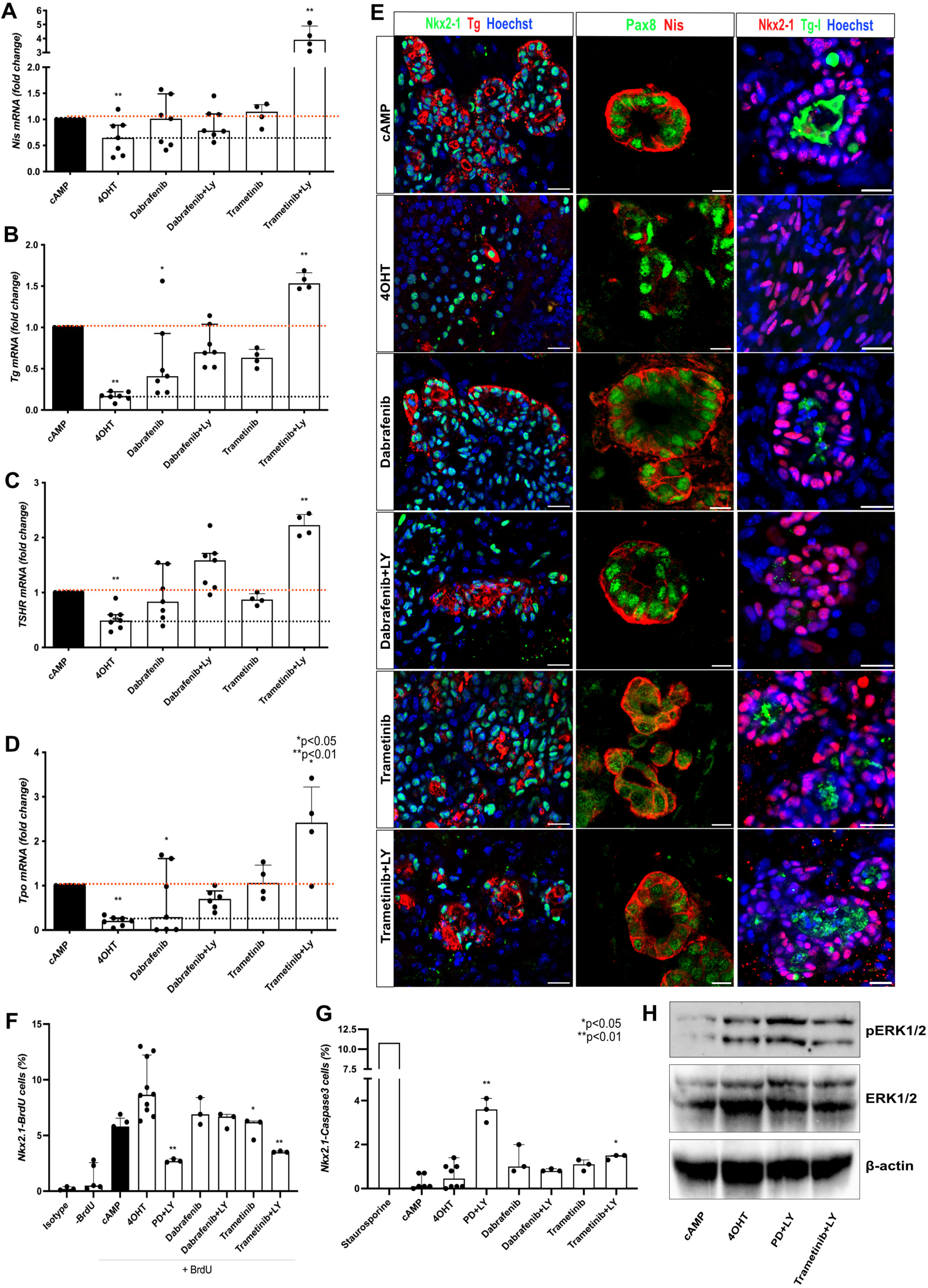
Dabrafenib and trametinib effect on redifferentiation of Braf^V637E-^expressing cells. Braf^V637E^-expressing organoids treated with Dabrafenib and Trametinib restore the expression of *Nis* to cAMP levels. However, a greater increase is observed under Trametinib+LY co-treatment (A). Recovery of *Tg* (B)*, TSHR* (C) and *Tpo* (D) mRNA to control levels was observed under co-treatment of Dabrafenib and Trametinib with the PI3K inhibitor (LY). Bar graphs represent the median (IQR) of at least 4 independent experiments with individual values shown. Comparisons were performed against the cAMP condition. (*p<0.05 ; **p<0.01; Mann-Whitney U test). Confocal images show that Dabrafenib and Trametinib isolated or combined to PI3K inhibitor (LY) also induce Tg and Nis protein levels while restoring the follicular structure (in a proportion of cells) and Tg-I accumulation in the lumen (E). Hoescht (shown in blue) was used for nuclei staining. Scale bars, 20 μm (Tg and Tg-I) and 10 μm (Nis). Proportions of proliferating (Nkx2.1/BrDU+) (F) and apoptotic (Nkx2.1/Caspase3+) (G) cells among control (cAMP), Braf^V637E^-induced (4OHT) and inhibitors (4OHT+inhibitors)-treated organoids after. For the proliferation assay, isotype and -BrdU conditions were used as negative controls for flow cytometry gating. As a positive control for apoptosis, staurosporine treatment (24 h) was used to determine Caspase 3 expression. Bar graphs represent the median (IQR) of at least 3 independent experiments with individual values shown. Comparisons were performed against the 4OHT condition. (*p<0.05 ; **p<0.01 ; Mann-Whitney U test). Western blot shows an increase of phospho-ERK (pERK) 7 days after the addition of 4OHT to the organoids culture when compared to cAMP control. Conversely, the treatment with Trametinib+LY resulted in pERK reduction compared to the 4OHT condition. (B). Beta-actin was used as a loading control for the immunoblot experiments. The image represents one experiment from 3 experimental replicates.

### Transcriptomic characterization of the PTC model and drug screening

Transcriptomic analysis of control (cAMP), Braf^V637E^-induced (4OHT) and inhibitors-treated (PD+LY) organoids (Fig. 6A) confirmed downregulation of thyroid genes under Braf^V637E^ stimulation (4OHT condition) whereas, global recovery of expression was observed under inhibitors treatment (Fig. 6A). Accordingly, thyroid differentiation scores (TDS and enhanced (e)TDS) were decreased under Braf^V637E^ induction while they recovered in the presence of inhibitors (Fig. 6B). Conversely, the ERK activation score was higher in the Braf^V637E^ samples and decreased sharply under inhibitor conditions (Fig. 6C). Since bulk RNAseq was performed using the follicle-enriched population, the presence of non-thyroid cells is reduced compared with the whole original organoids but still present. It may explain the modest increase in ERK score among Braf^V637E^-activated cells while the apparent decrease under the condition with inhibitors reflects the effect of the treatment on each cell type.

**Fig. 6.**
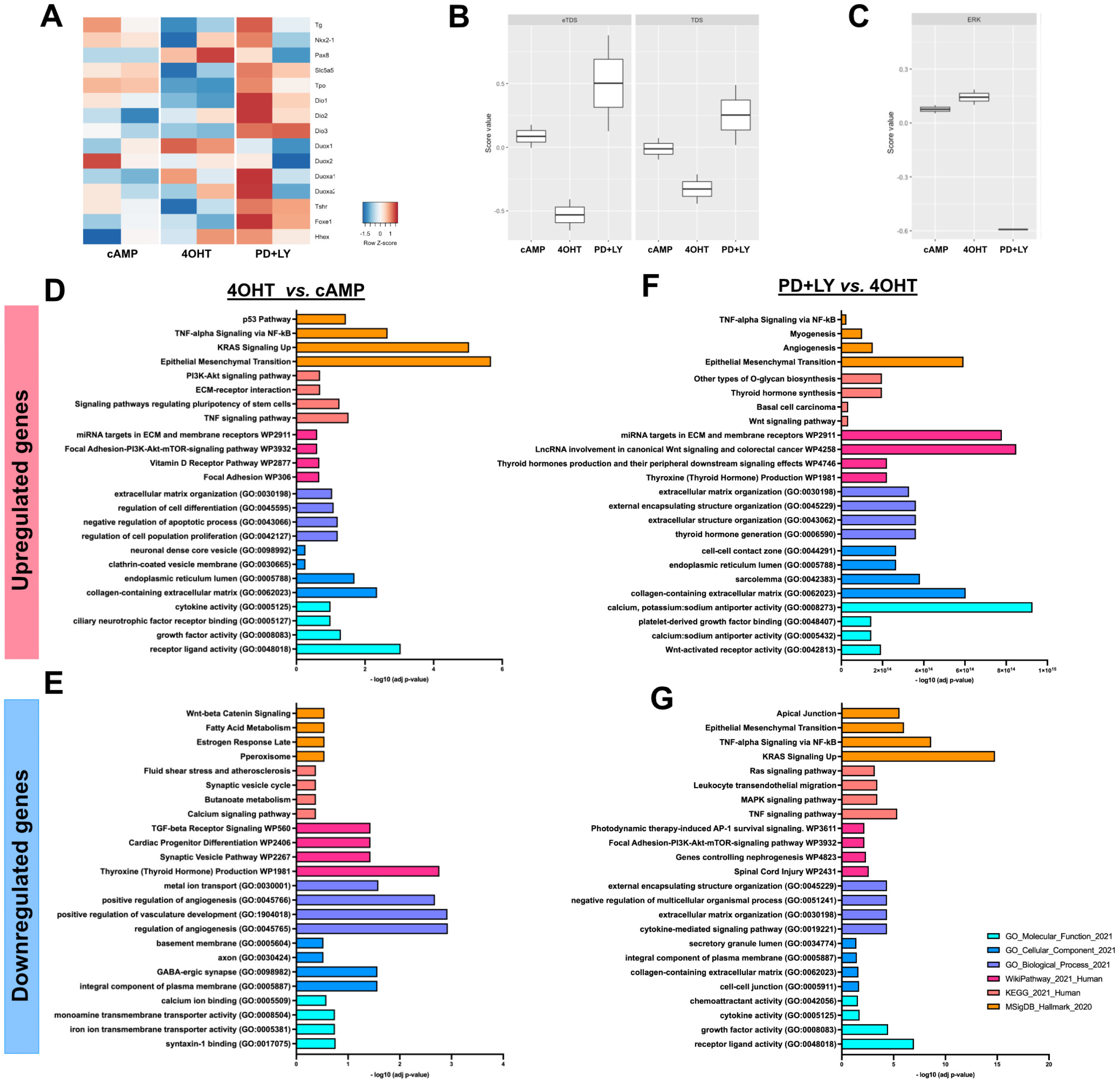
Transcriptomics analysis confirms thyroid redifferentiation of Braf^V637E^- expressing cells treated with MAPK and PI3K inhibitors and suggests by which mechanisms. Heatmap of normalized bulk RNA-Seq expression of normal thyroid cells (cAMP), Braf^V637E^ expressing cells (4OHT), and Braf^V637E^ expressing cells treated with PD0325901+LY204002 inhibitors. Rows represent markers and columns represent specific conditions. Color values in the heatmap represent mean expression levels (A). Thyroid differentiation (eTDS and TDS) (B) and ERK (C) scores were calculated among the different conditions. Classification of upregulated and downregulated genes comparing 4OHT *vs.* cAMP (D, E) and PD+LY *vs.* 4OHT (F, G). Colors represent the classification method and scale the -log10 (adj p-value).

Differential expression analysis identified 321 Differential Expressed Genes (DEGs; 156 upregulated and 165 downregulated genes) in the 4OHT condition compared to cAMP; and 853 DEGs in inhibitors compared to 4OHT (287 upregulated and 566 downregulated genes). Gene enrichment classification analysis of DEGs results is presented in Fig. 6D-G. Briefly, among the upregulated genes in Braf^V637E^-induced compared to control, we observed gene signature for the hyperactivation of PI3K-AKT-mTOR, TNF, and cytokines signaling and promotion of Epithelial-Mesenchymal transition (EMT) (Fig. 6D). While genes associated with thyroid hormone production, TGF-beta, Wnt/Beta-Catenin pathway, and regulation of angiogenesis were down-regulated (Fig. 6E). Conversely, the inhibitor’s treatment, compared to 4OHT condition, evidenced upregulation of genes associated with thyroid hormone production, cell-cell contact, extra-cellular matrix organization, and angiogenesis processes (Fig. 6F), while PI3K, MAPK, TNF, cytokines signaling, ECM-receptor interactions and EMT-related genes were downregulated (Fig. 6G). The list of the DEGs for each gene enrichment classification is provided in Supplementary Table 4.

Given our transcriptomics analysis, which indicated elevated expression of inflammation-related genes and downregulation of the Wnt/β-catenin pathway in Braf^V637E^-expressing cells, we tested the effect of anti-inflammatory drugs, specifically dexamethasone and CC-5013 (Lenalidomide; TNF-alpha inhibitor) as well as CHIR-99021 (Wnt/β-catenin activator) on our cancer organoids. Following the same experimental strategy as with the other inhibitors (Fig. 4A), we observed that co-treatment with dexamethasone and CC-5013 led to approximately a 3.5 fold increase in *Nis* expression compared to the control condition (cAMP) (Supplementary Fig. 9A), confirming the previously described inhibitory effect of inflammation on *Nis* expression. On the other hand, for the other main thyroid differentiation genes, namely *Tg, TSHR* and *Tpo*, treatment with anti-inflammatory drugs and the Wnt/β-catenin activator did not result in their re-expression (Supplementary Fig. 9B-D). This suggests that these alterations are likely a consequence of the oncogenic process rather than being driving factors.

## DISCUSSION

Here, we described a Braf^V637E^ oncogene-derived thyroid cancer organoid model recapitulating patient tumor features. *In vitro* thyroid cancer models that recapitulate tumor development and behavior can facilitate the identification of early tumor drivers and enable the screening of several new drugs for the treatment of thyroid cancer without the need to use too many animal models. Combining our previous thyroid organoid model derived from mESCs with an inducible system, the Braf^V637E^ oncogene could be induced explicitly in mature Tg-expressing cells to obtain a 3D *in vitro* cancer model. Overexpression of Braf^V637E^ rapidly led to MAPK activation with increased pERK, cell dedifferentiation, and disruption of follicular organization. Similar effects have been described in genetically modified mouse models^41,59,73^. Furthermore, the gene expression signature of our cancer organoids confirmed the findings found in the PTC samples, which also showed enrichment of genes associated to p53, focal adhesion, ECM-receptor interactions, EMT and inflammatory pathways^74–76^.

Activating mutations in the *BRAF* gene are found in approximately 7% of all solid human tumors, particularly common in PTCs, ATCs, and melanomas. In addition, they have been reported less frequently in colorectal cancers (CRCs), lung cancers, pediatric low-grade gliomas (PLGGs), glioblastomas, breast cancers, and serous ovarian cancers^77–83^. In PTCs and melanomas, BRAF^V600E^ mutation seems to be associated with a higher degree of dedifferentiation and more aggressive histological patterns. However, its prognostic role is still debated as it was not found independently of histological features^46,47^. Nevertheless, studies suggest that BRAF^V600E^ might predict response to tyrosine kinase inhibitors (TKIs) in melanoma and lung cancer^84,85^.

Surgery remains the first choice for thyroid cancer therapy. Recommended post-operative treatment includes TSH suppression and RAI ablation, particularly as adjuvant treatment for patients at high risk for tumor recurrence and to treat patients with persistent/recurrent or metastatic disease^86^. The benefit of RAI has been demonstrated in patients by reducing the risk of recurrence and disease-related mortality^87^. However, approximately two-thirds of metastatic DTC become radioiodine refractory (RR-DTC), defined by the absence of iodine uptake or tumor progression despite uptake^88,89^. There is no curative treatment for radioiodine refractory DTCs. The recommended first-line systemic treatment when local therapies are not possible is targeted therapies using mainly multitarget TKIs, such as Lenvatinib showed a progression-free survival of 18 months ^67,90–92^.

NIS, a basal membrane iodide transporter, plays a critical role in radioiodine accumulation in DTC cells and its level is closely related to response to RAI (I^131^) therapy. Studies in mouse models and humans have shown that the presence of *BRAF* mutations results in lower *Nis* levels and, consequently a higher rate of RAI-refractory tumours^41,44^. Here, we confirmed that this regulation is preserved in our cancer organoids, as disruption of Braf^V637E^ activation led to in recovery of *Nis* levels. As TKIs, retinoic acids (RA), histone deacetylase inhibitors (HDAC), peroxisome proliferator-activated receptor-gamma (PPARG) have been tested to promote redifferentiation and *NIS* re-expression of RR-DTCs, to suggest RAI treatment after a short, targeted treatment^90,93–95^. This strategy could lead to tumor response while limiting adverse effects and, several clinical trials are ongoing. However, the re-expression of NIS is insufficient to explain the response to redifferentiation therapeutics and RAI treatment. Membrane trafficking and the cellular machinery that concentrate and retain iodine must be preserved^96^ in a follicular organization. Our model is proving to be a potential tool for redifferentiation studies because of its ability to test large sets and combinations of treatments and to assess follicular reorganization and iodide organification capacity using a functional assay that is likely to be more strongly associated with response to RAI treatment than *NIS* mRNA levels. Braf^V637E^-expressing cells treated with VPA, MAPK, PI3K, Dabrafenib, Trametinib, and inflammation inhibitors were shown to restore the expression of *Nis*. Interestingly, the combination of MAPK (BRAF and MEK) and PI3K inhibitors also promoted the restoration of thyroid markers, follicular organization, and iodide organification ability. Interestingly, an ongoing clinical trial tests the effect of BKM120, a PI3K inhibitor, in patients with progressive, metastatic, refractory, follicular or poorly differentiated thyroid cancers (NCT01830504). However, the association with MAPK inhibitors is still not evaluated in patients with thyroid cancer.

In *BRAF*-mutated tumors, studies have shown that the combination of TKIs therapies appears more effective and circumvents primary and acquired resistance to TKI therapy. Often, resistance is due to reactivation of the MAPK/ERK pathway or activation of other signaling pathways such as PTEN, NF-1 or RAS. It may also result from the hyperactivation of tyrosine kinase receptors, such as PDFRβ, IGF-1R and HGF, which lead to activation of the AKT/PI3K pathway^97^. Studies in patients with advanced melanoma carrying a BRAF^V600E^ mutation have shown that combining BRAF and MEK inhibitors resulted in a higher rate of complete/partial responses and median progression-free survival compared with monotherapy groups^98–100^. Such significant results lead to the use of drug combinations as standard treatment for patients^101,102^. Likewise, phase II clinical trials have demonstrated noteworthy response rates in advanced *BRAF*-mutated thyroid carcinomas when treated with a combination of the BRAF inhibitor dabrafenib and MEK inhibitor trametinib^70^. Furthermore, recent findings have shown that the administration of dabrafenib-trametinib treatment followed by surgery can yield 24-month overall survival (OS) rates as high as 80% in ATCs^103^.

In summary, we have developed a Braf^V637E^ oncogene-expressing thyroid cancer organoid *in vitro* model from mESCs that recapitulates transcriptomic and histological features of PTCs at early and advanced stages. Moreover, we demonstrated that the combination of MEK and PI3K inhibitors promotes *Nis* re-expression and cell re-differentiation leading to the restoration of follicular structures and thyroid functionality. Considering the robustness of this *in vitro* model, which allows controlled induction of the major thyroid oncogene in a three-dimensional system, its efficiency and simplicity, our model can be used to study mechanisms associated with thyroid cancer development and progression, thyroid redifferentiation, and drug screening.

## MATERIAL AND METHODS

### Cell culture and mESC_Braf^V637E^ line generation

The previously genetically modified G4 RosaLuc mouse embryonic stem cell line (mESC)^104^ was initially cultured on gamma-irradiated mouse embryonic fibroblasts (MEFs) feeder using mouse stem cell medium^4,58^ and incubated at 37 °C, 5 % CO^2^ and >95 % humidity. Cells were routinely tested for mycoplasma. To insert the target sequences into the modified Rosa26 locus of G4 RosaLuc mESCs, around 1×10^5^ cells were transfected using the Flpe expressing vector^104^ and the target vector containing the rtTA-TRE induction system, thyroid transcription factors, Nkx2-1 and Pax8 and bTg-NES-Braf^V637E^-ERT^2^ sequences (Supplementary Fig. 1A) following the lipofectamine 3000 protocol (Thermo Scientific). The Braf^V637E^ mutation here used for mouse cells is equivalent to the human BRAF^V600E^ mutation^56^. Briefly, 1×10^6^ mESCs were transfected (in suspension) with 20 μg of each vector and cultured on Neomycin-resistant MEFs. G418 (Neomycin; 300 μg/ml) was applied 48 h after transfection, for 10 days, and individual clones were isolated (colonies were separated from the MEFs using insulin needles) and further expanded. Clones were screened by immunofluorescence for Nkx2-1 and Pax8 expression after three days of incubation with 1 µg/ml Doxycyclin (Dox) (Supplementary Fig. 1B). Positive clones were then characterized according to pluripotency maintenance (Supplementary Fig. 1C), *Nkx2-1* and *Pax8* expression, and efficiency of thyroid differentiation. In addition, we generated an mESC control line where the sequences of NES-BRAF^V637E^-ERT^2^ were replaced by the eGFP sequence, resulting in the TRE-Nkx2-1-Pax8_bTg-eGFP line (Supplementary Fig. 1D). The mESC control line generation, selection and characterization were performed as described above (Supplementary Fig. 1E, F).

### Thyroid differentiation protocol

G4 RosaLuc TRE-Nkx2-1-Pax8_bTg-NES-Braf^V637E^-ERT^2^ and TRE-Nkx2-1-Pax8_bTg-eGFP cells were cultured and differentiated as described previously by Antonica *et al.* (2012)^4^ with few modifications. Briefly, modified mESCs cultured in mESC media on top of MEFs were split using Trypsin EDTA (TE). Then, cells were resuspended in mESC media^57^ and seeded into a 10-cm petri dish for 30-45 min allowing the attachment of most of the MEFs. mESC-enriched supernatant was collected, centrifuged (500g for 5min), and resuspended in differentiation media^57^, cells were counted and finally diluted to 40,000 cells/ml. Embryoid bodies (EBs), were then generated by hanging drops (25µl drops containing 1,000 cells), collected after four days and embedded in growth factor reduced Matrigel (GFR MTG; BD Biosciences); 50µl MTG drops (containing around 20 EBs) were plated into 12-wells plates. EBs were differentiated using a differentiation medium (1 ml/well) initially supplemented with 1µg/ml of Doxycycline (Sigma) for five days, followed by two weeks of maturation induction by using 0.3 µmol of 8-Br-cAMP (BioLog Life Science Institute) or 1mU ml^-1^ of thyrotropin (hrTSH; Genzyme). The culture medium was changed every 2 days. Thyroid differentiation and functionality were evaluated by transcriptomics (RT-PCR and bulk RNA sequencing), immunofluorescence, and iodide organification.

### Braf^V637E^ induction and drug screening

After full thyroid maturation (Day 23), MTG drops were washed twice with Hanks’s balanced salt solution (HBSS, containing calcium and magnesium; Invitrogen) and incubated in a HBSS solution (1 ml per well) containing 10 U/ml of dispase II (Roche) and 125 U/ml of collagenase type IV (Sigma) for 30 min at 37°C. Then the enzymes were inactivated by adding 10% FBS and cells were centrifuged at 500g for 3 min. Cells were rinsed twice with HBSS, and the follicle population was enriched by filtering using 30 μm (single cell removal) and 100 μm reverse strainer (Pluriselect). Finally, enriched structures were re-embedded in GFR MTG (50µl) and plated into 12-well plates. Twenty-four hours later, cells were incubated with 1 μM of 4-Hydroxytamoxifen (4OHT) (Sigma) and 8-Br-cAMP or hrTSH for 24h, 48h,7 days or 3 weeks to promote Braf^V637E^-induced phosphorylation of MAPK pathway^56^. The control condition, treated with cAMP and/or hrTSH, was also treated with vehicle ethanol for 4OHT treatment comparisons.

Braf^V637E^-induced thyroid organoids (previously treated for four days with 4OHT) were also cultured in addition to distinct compounds, previously suggested to regulate Nis re-expression and inhibit Braf^V637E^ oncogenic effects^68,69^. Among the screened drugs, several cell processes were targeted by the following compounds: PD0325901 (MEK inhibitor; 250nM; Stem Cell), LY204002 (PI3K inhibitor; 5μM; Selleckchem), VPA (HDAC inhibitor; 250μM; Sigma), NAC (N-acetyl cysteine, antioxidant compound; 2mM; Sigma), 5-AZA-2’ deoxycytidine (DNA methyltransferase (DMNT) inhibitor; 1μM; Sigma) and RG108 (DMNT inhibitor; 10μM; Sigma). Furthermore, experiments were also conducted using inhibitors that are already clinically approved for the treatment of thyroid cancer, namely dabrafenib (BRAF inhibitor; 100nM; Selleckchem) and/or trametinib (MEK inhibitor; 20nM; Selleckchem).

Disrupted pathways identified by transcriptomics analysis in 4OHT condition compared to cAMP control were inhibited/activated by treatment with dexamethasone (anti-inflammatory drug; 50nM; Sigma), CC-5013 (Lenalidomide; TNF-alpha inhibitor; 10μM; Selleckchem) and CHIR-99021(Wnt/β-catenin activator; 3μM; Selleckchem). All inhibitor treatments involved organoids that had previously been exposed to cAMP+4OHT for four days, and they were subsequently subjected to continuous incubation with cAMP+4OHT+inhibitors for an additional 3 days. Culture media was replenished every 2 days. All experiments were performed following the same time points of incubation described above.

### Flow cytometry

#### Proliferation assay

Organoids subjected to various treatments, including controls (cAMP), 4OHT, and 4OHT+inhibitors, at both 7 days (day 30) and 21 days (day 54), were exposed to BrdU (1mM) for a duration of 3 hours. Subsequently, a proliferation assay was conducted in accordance with the BrdU Flow Kit staining protocol (BD) instructions. Briefly, Matrigel drops (pool of at least four replicates per condition) were first digested with HBSS solution containing 10 U/ml dispase II and 125 U/ml collagenase type IV for 30-60 min at 37°C; then a single cell suspension was obtained by dissociation with TripLE Express (10-15 min incubation at 37°C), the enzymes were inactivated by addition of differentiation medium. After centrifugation (500 g for 3 min), samples were rinsed with PBS, and then the BrdU Flow Kit staining protocol (BD Biosciences) was used according to the manufacturer’s instructions. In order to identify the BrdU incorporated cells among the thyroid population, cells were also stained using primary Nkx2-1 antibody (1:100; Abcam) and donkey anti-rabbit IgG Cy3-conjugated (1:300; Jackson Immunoresearch). As controls, we used cells untreated with BrdU and stained with the APC anti-BrdU antibody for BrdU gating while for Nkx2.1 gating, cells were incubated only with the secondary antibody. Data are presented as a percentage of double-positive (BrdU/Nkx2.1) cells. For each experiment, at least four wells from each condition were used.

#### Apoptosis assay

Organoids subjected to various treatments, including controls (cAMP), 4OHT, and 4OHT+inhibitors, at both 7 days (day 30) and 21 days (day 54) were collected and dissociated into single cells following the protocol described above (proliferation assay section). Cells were then fixed and permeabilized using the BD Cytofix/Cytoperm™ Fixation/Permeabilization Kit (BD Biosciences) and stained using Nkx2-1 antibody (1:100; Abcam) and donkey anti-rabbit IgG Cy3-conjugated secondary antibody (1:300; Jackson Immunoresearch) combined with FITC Rabbit Anti-Active Caspase-3 conjugated antibody (1:5; BD Biosciences). As a positive control, cAMP-treated organoids were incubated with Staurosporine (10nM; Sigma) for 16 hours and collected at the respective time points following the procedures described above. Untreated cells and cells incubated only with the secondary antibody (isotype) were used as negative controls for the gating strategy. For each experiment, at least four wells from each condition were used.

#### Proportion of bTg-GFP cells

The proportions of Nkx2.1 and GFP cells in the control mESC line (TRE-Nkx2-1-Pax8_bTg-eGFP) were evaluated at day 30. Controls (cAMP) and 4OHT-treated (7 days) organoids were dissociated and stained for Nkx2.1 following the steps described above (apoptosis assay session). GFP detection was based on the endogenous expression (bTg-eGFP). Undifferentiated cells (mESCs) and cells incubated only with the secondary antibody (isotype) were used as negative controls for the gating strategy. For each experiment, at least four wells from each condition were used.

All analyses were performed using the LSR-Fortessa X-20 flow cytometer and BD FACSDiva software was used for the quantification analysis. Data are presented as a percentage of positive cells.

### Immunofluorescence staining

Organoids embedded in MTG were fixed in 4% paraformaldehyde (PFA; Sigma) for 1 h at room temperature (RT) and washed three times in PBS. Samples were either stained as whole mount (MTG drop) or embedded in 4% low-melting agarose and cut using Vibratome (80-100μm; Sigma). Blocking was performed using a solution of PBS containing 3% bovine serum albumin (BSA; Sigma), 5% horse serum (Invitrogen) and 0.3% Triton X-100 (Sigma) for 30 min at RT. The primary and secondary antibodies were diluted in a solution of PBS containing 3% BSA, 1% horse serum and 0.1% Triton X-100. Primary antibodies were incubated overnight at 4°C followed by incubation with secondary antibodies and Hoechst for 2 h at RT. Slides were mounted with Glycergel (Dako).

For paraffin embedding, organoids were fixed in 4% PFA for 1 h at 4°C and kept in 70% ethanol at 4°C before processing. Samples were then embedded in paraffin, sectioned (5 μm), mounted on glass slides, deparaffinized, and rehydrated following standard protocols. For immunostaining, antigen retrieval was performed by incubating the sections for 10 min in the microwave (850 W) in Sodium Citrate Buffer (10 mM Sodium Citrate, 0.05% Tween 20, pH 6.0). After cooling, the sections were rinsed with PBS and blocked with 1% BSA and 10% horse serum PBS solution for 1 h at RT. Primary antibodies (Supplementary Table 1) were diluted in the blocking solution and incubated overnight at 4°C. The sections were rinsed three times in PBS and incubated with Hoechst and secondary antibodies diluted in blocking solution for 1 h at RT. Slides were mounted with Glycergel (Dako). Imaging was performed using a Zeiss LSM510 META confocal microscope and a Leica DMI6000 microscope with DFC365FX camera. The antibodies specifications and dilutions are listed in the supplementary Table 1.

### Western blot

Follicle enriched-organoids were initially isolated (collagenase IV/dispase II solution, described above) from at least 6 distinct wells and pooled together for protein extraction using RIPA buffer. The protein concentration was determined according to Pierce 660 nm protein assay reagent protocol (Thermo Scientific). Briefly, for each sample, 30–50 μg of protein was fractionated by 10% SDS–PAGE and blotted onto an Immobilon PVDF membrane (Millipore). Non-specific binding sites were blocked by incubation with 5% BSA in Tris-buffered saline (TBS) 0.1% Tween-20 (TBS-T) for 1 h. Thereafter, the membrane was incubated overnight at 4 °C with primary antibodies against phospho-ERK1/2 (1:1,000; Cell Signaling), ERK1/2 (1:400; Santa Cruz Biotechnology), and β-Actin (1:1,1000; Cell Signaling) in a blocking solution. Next, the membrane was incubated with horseradish peroxidase-conjugated anti-rabbit (1:1,000; Cell Signaling) or anti-mouse antibody (1:1,000; Cell Signaling) in a blocking solution, for 60 min at RT. The antigen-antibody complexes were visualized using an enhanced chemiluminescence system (GE Healthcare) and captured by Azure 500 system (Azure Biosystems).

### Iodide organification assay

Thyroid organoids treated with cAMP, cAMP+4OHT, cAMP+4OHT+PD+LY and cAMP+4OHT+PD+LY+VPA were tested for the ability of iodide uptake and organification as previously described^4,58^. Briefly, cells were washed with HBSS and incubated with 1 ml per well of an organification medium containing 1,000,000 c.p.m. per ml ^125^I (PerkinElmer) and 100 nM sodium iodide (NaI, Sigma) in HBSS for 2 h at 37 °C. The reaction was stopped by the addition of 1 ml of methimazole (MMI; 4 mM, Sigma). After two washes with cold PBS, organoids were dissociated by incubation with trypsin/EDTA (Invitrogen) for 10 min at 37 °C. For iodide uptake quantification, cells were collected, and radioactivity was measured using a gamma-counter. Subsequently, proteins were precipitated by adding 100 μl of gamma-globulins (10 mg/ml; Sigma) and 2 ml of 20% TCA followed by centrifugation at 2,000 r.p.m. for 10 min, at 4°C and the radioactivity of protein-bound ^125^I (PBI) was measured. Iodide organification was calculated as an iodide uptake/PBI ratio and, the values were expressed as a percentage. Cells were also treated with 1 mM sodium perchlorate (Nis inhibitor; NaClO4, Sigma-Aldrich) and 2 mM MMI (Tpo inhibitor; Sigma-Aldrich) as iodide uptake and protein-binding controls, respectively. The experiments were performed in triplicates for each condition.

### Gene expression analysis

Real-time PCR (RT-qPCR) was performed on cDNA from thyroid organoids from at least three independent experiments. Total RNA was extracted from thyroid organoids by the addition of lysis RLT Lysis buffer (Qiagen) + 1% 2-mercaptoethanol directly on the MTG drop containing the organoids. For longer cultures (21 days), organoids were initially incubated with a collagenase IV/Dispase II solution (described above) for 30-45 min. Enzymes were inactivated by the addition of differentiation medium, cells were centrifuged (500 g for 3 min), washed with PBS, and resuspended in RLT buffer. RNA was isolated using the RNeasy micro kit (Qiagen) according to the manufacturer’s instructions. cDNA was generated by reverse transcription using the Superscript II kit (Invitrogen). qPCR was performed in triplicates on cDNA (1:10 dilution) using Takyon™ No ROX SYBR 2X MasterMix blue dTTP (Eurogentec) and CFX Connect Real-Time System (Biorad). Results are presented as linearized values normalized to the housekeeping gene, β2-microglobulin and the indicated reference value (2-DDCt). cAMP condition was used as the control for all the comparisons. Primer sequences are described in Supplementary Table 2.

### Bulk RNA sequencing and analysis

Bulk RNA-seq was performed on day 31, using the cAMP, cAMP+4OHT (5 days) and cAMP+4OHT+PD+LY (3 days) conditions (see protocol, Fig. 5A). RNA extraction was performed as previously described (section “*Gene expression analysis”*), and experiments were performed in duplicates. At least three distinct wells were pooled together for each condition. RNA quality and concentration were evaluated using Bioanalyser 2100 (Agilent) and RNA 6000 Nano Kit (Agilent). RNA integrity was preserved, and no genomic DNA contamination was detected. Indexed cDNA libraries were obtained using the TruSeq Stranded mRNA Sample Prep kit (Illumina) using an S2 flow cell, and sequences were produced using a 200 cycles reagent kit. The resulting high-quality indexed cDNA libraries were quantified with the Quant-iT PicoGreen kit (Life Sciences) and Infinite F200 Proplate reader (Tecan); DNA fragment size distribution was examined with the 2100 Bioanalyzer (Agilent) using the DNA 1000 kit (Agilent). Multiplexed libraries (10ρM) were loaded onto flow cells and sequenced on the HiSeq 1500 system (Illumina) in high-output mode using the HiSeq Cluster Kit v4 (Illumina). The sequenced data were uploaded on the galaxy web platform version 22.05.1, and the public server, Data independent acquisition proteomics workbench (RRID:SCR_021862, https://usegalaxy.eu), was used for mapping and counting analysis. Approximately 10 million paired-end reads were obtained per sample. After the removal of low-quality bases and Illumina adapter sequences using Trimmomatic software^105^ (RRID:SCR_011848), paired-end reads were mapped against the mouse reference genome mm10 (GRCm38) using HISAT2 sofware^106^, allowing soft clipping. Raw read counts were obtained using HTseq count software with unstranded option^107^. Low-expressed genes were filtered using the EdgeR package in R Project for Statistical Computing (https://www.r-project.org). Then, differential expression analysis was performed using DESeq2 package^108^. Fold change and adjusted p-value thresholds of 2 and 0.05, respectively were used to select differentially expressed genes (DEG). Gene ontology and pathways enrichment analysis in up and down differentially expressed gene lists was realized using EnrichR (https://maayanlab.cloud/Enrichr/). Transformed counts corresponding to log2 (normalized counts +4) using the DESeq2 package were used for further analysis and heatmap visualization. Thyroid differentiation score (TDS), Enhanced Thyroid differentiation score (eTDS) and ERK activation score were calculated as previously described^67^. The list of the genes used for the score’s calculation is provided in Supplementary Table 3.

### Data availability

Bulk RNA-seq data have been deposited in the NCBI Gene Expression Omnibus (GEO; RRID:SCR_005012) under accession number GSE228281.

### Statistical analysis

All statistical analysis were performed using GraphPad Prism 9. Comparison between two groups and its statistical significance was tested using the nonparametric Mann-Whitney U test. In contrast, comparisons between multiple groups was performed using the nonparametric Kruskal-Wallis test. Data are displayed as median (IQR). Differences were considered significant at p<0.05 and presented as follows: *p < 0.05, **p < 0.01, ***p < 0.001, ****p < 0.0001. All the data presented are from at least three independent experiments.

## Acknowledgements and Funding

The authors acknowledge the ULB flow cytometry platform (C. Dubois), the ULB genomic core facility (F. Libert and A. Lefort), LiMIF platform for confocal microscopy (J.-M. Vanderwinden) and Eduardo Andrés Rios Morris for the contribution with the imaging. We acknowledge the funding agencies that supported this study. The Belgian National Fund for Scientific Research (FNRS) (PDR T.0140.14; PDR T.0230.18, CDR J.0068.22, Televie 7.4633.17/7.4526.19) and the Fonds d’Encouragement à la Recherche de l’Université Libre de Bruxelles (FER-ULB). The Belgian Fondation contre le cancer (F/2020/1402; V.D.). FNRS (Chargé de Recherche, No.825745; M.R.). The cooperation program CAPES-WBI (S.C. and A.L.M). The Brazilian National Council for Scientific and Technological Development (CNPq; M.R. and A.L.M), the Coordination for the Improvement of Higher Education Personnel (CAPES; M.R. and A.L.M), the Brazilian Society of Endocrinology and Metabolism (SBEM; M.R. and A.L.M) and the Fundação de Amparo à Pesquisa do Estado do Rio Grande do Sul (FAPERGS; A.L.M). S.C is Research Director at FNRS. M.R is researcher at ULB.

## Author contributions

H.L., S.C. and M.R. developed the project, designed the experiments and analyzed the data. L.H., T.P., S.G. and J.H. generated the plasmids and the G4 Rosaluc mESC line. M.R. and A.S. generated the G4 RosaLuc TRE-Nkx2-1-Pax8_bTg-NES-Braf^V637E^-ERT^2^ and TRE-Nkx2-1-Pax8_bTg-eGFP cell lines. H.L., M.R., B.F.F., A.S., O.M., L.C., M.K.P. B.A., L.C., performed the *in vitro* experiments and protocol set up. M.R. and P.G. obtained confocal images. H.L., M.R. and A.T. performed bulk RNA-Sequencing and analyzed the results. H.L. and A.T. performed the bioinformatics analysis. H.L. and M.R. wrote the first draft and, A.L.M. and S.C. edited the manuscript. S.C., M.R. and A.L.M. acquired funding for the project. All authors contributed to the article and approved the submitted version.

**Correspondence** and requests for materials should be addressed to Mírian Romitti.

