## Supplementary figures and tables for "Dual targeting of MAPK and PI3K pathways unlocks redifferentiation of *Braf*-mutated thyroid cancer organoids"

### SUPPLEMENTARY MATERIAL

#### Supplementary figures

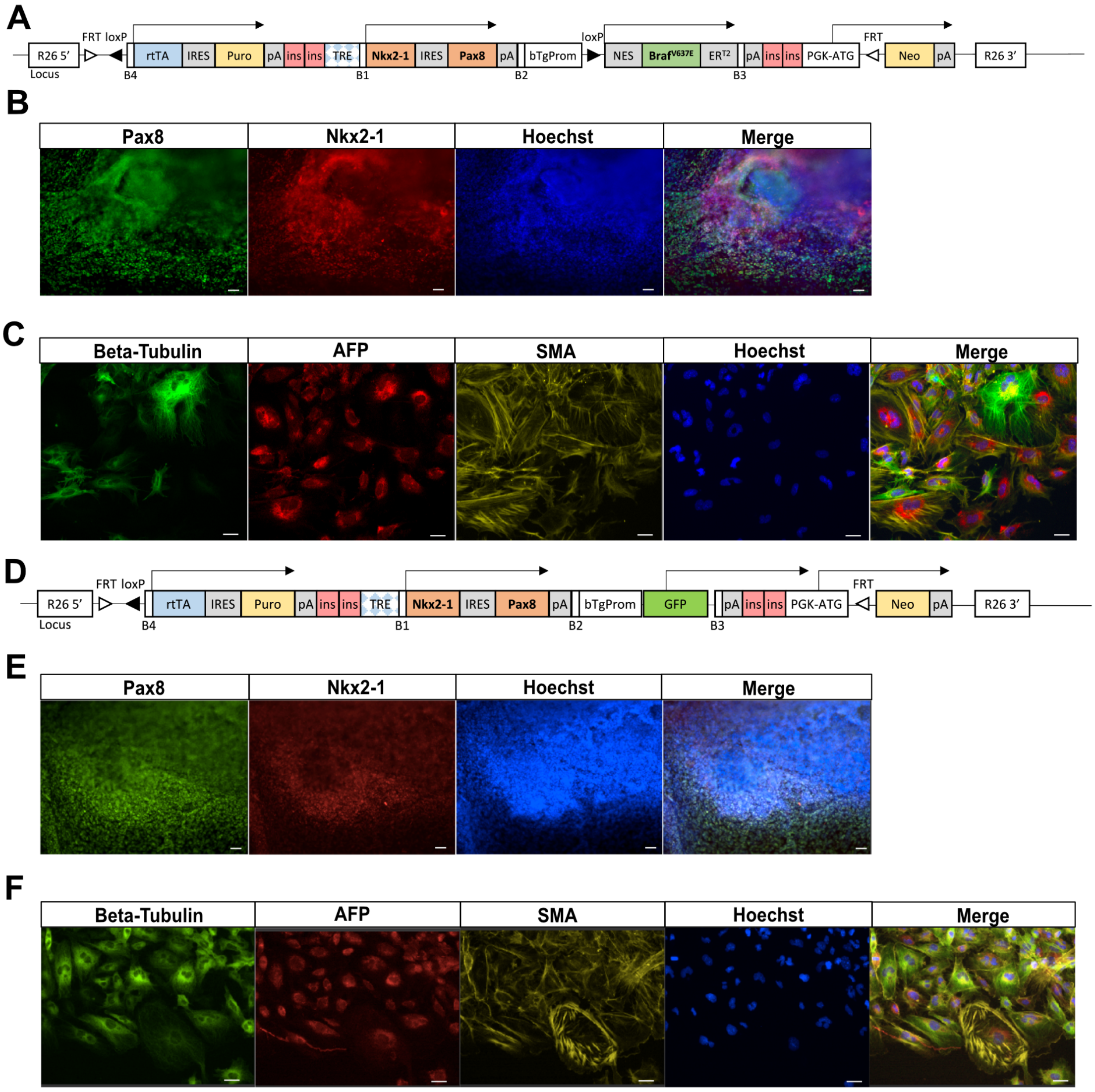


**Supplementary Fig. 1 mESC-cell lines generation and characterization.** Schematic representation of the mESC_Nkx2-1/Pax8_bTg_$Braf^{V637E}$ cell line (A). mESC_Nkx2-1/Pax8_bTg_$Braf^{V637E}$ line response to doxycycline treatment shows expression of Pax8 and Nkx2-1 proteins (B). Spontaneous differentiation mESC_Nkx2-1/Pax8_bTg_$Braf^{V637E}$ cell line leads to the generation of ectodermal (Beta-Tubulin), endodermal (AFP) and mesodermal (SMA) cells (C). Schematic representation of the mESC_Nkx2-1/Pax8_bTg_eGFP control cell line (D). Dox treatment leads to an increase in Pax8 and Nkx2-1 expression among control cells (E). Spontaneous differentiation of the control cells leads to the generation of ectodermal (Beta-Tubulin), endodermal (AFP) and mesodermal (SMA) cells (F). Scale bars, 50 µm (B, E) and 10 µm (C, F).

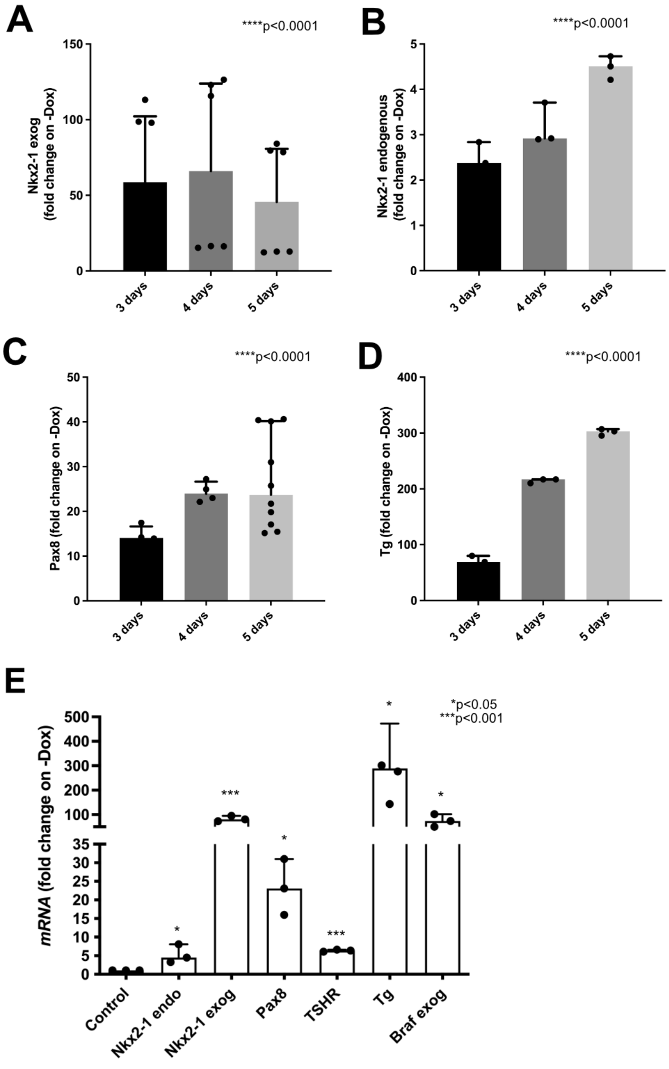


**Supplementary Fig. 2 Doxycycline treatment for 5 days leads to more efficient thyroid gene expression induction.** Time course of Dox-dependent induction of exogenous and endogenous *Nkx2-1* (A, B) *Pax8* (C) and *Tg* (D) expression. mRNA levels of thyroid genes at day 9, after 5 days of Dox treatment (E). Bar graphs represent the median (IQR) of at least 3 independent experiments with individual values shown.

Comparisons were performed against the -Dox condition. (*p<0.05 ; **p<0.01 ; ***p<0.001; ***p<0.0001; Kruskal-Wallis test (A-D) and Mann-Whitney U test (E)).

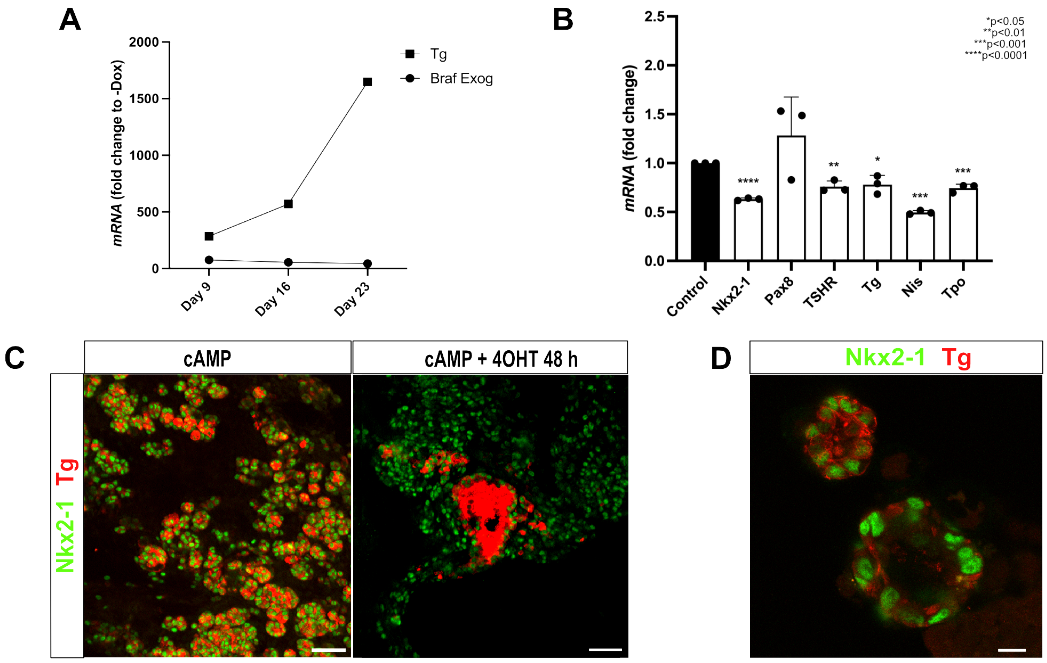


**Supplementary Fig. 3 4OHT treatment rapidly induces Braf-carcinogenic changes in gene expression and follicular organization.** Endogenous Tg and exogenous Braf expression levels at day 9, 16, and 23 of thyroid differentiation protocol (A). Induction of $Braf^{V637E}$ for 8 hours promotes the downregulation of main thyroid genes (B). Bar graphs represent the median (IQR) of 3 independent experiments with individual values shown. Comparisons were performed against the cAMP condition. (*p<0.05 ; **p<0.01 ; ***p<0.001; ***p<0.0001; Mann-Whitney U test). Nkx2-1 and Tg immunostaining shows that under $Braf^{V637E}$ activation by 4OHT (48 h) thyroid follicles present a bigger size and heterogeneous distribution of Tg compared to the control (cAMP) (C). Remaining follicular-like structures after $Braf^{V637E}$ activation (4OHT; 48 h) (D). Scale bars, 50 μm (C) and 10 μm (D).

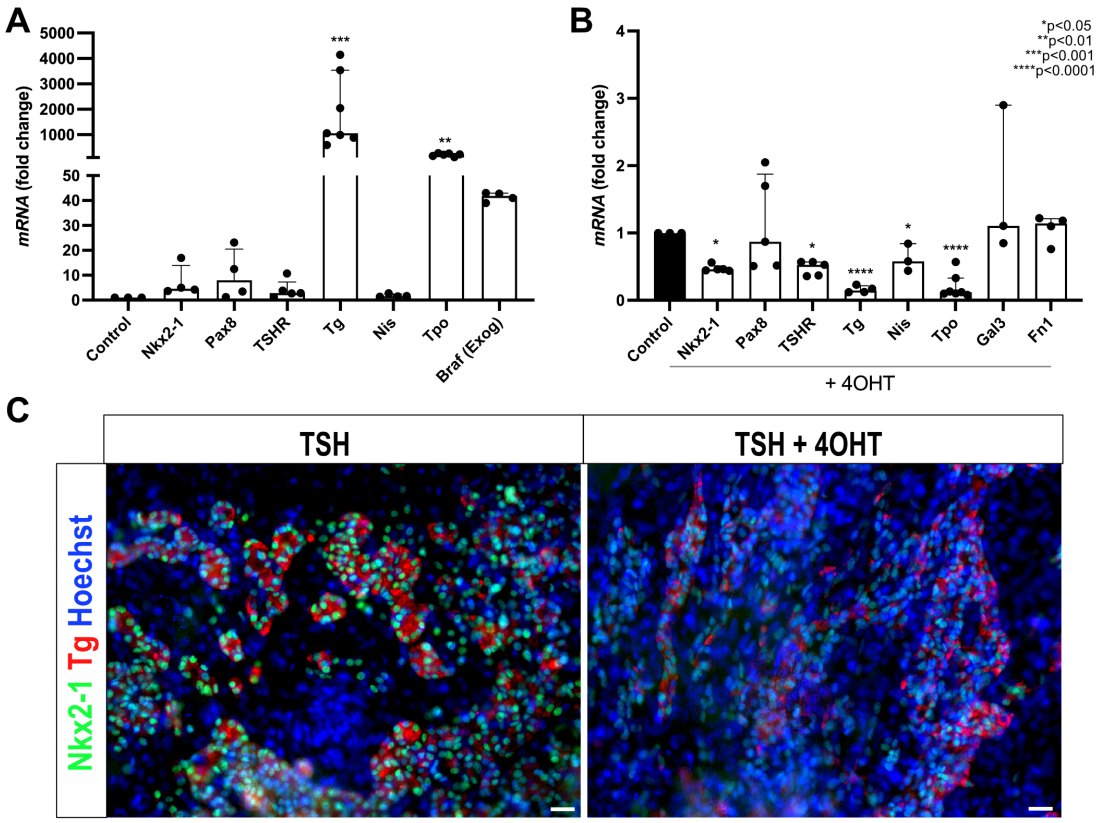


**Supplementary Fig. 4 Braf$^{V637E}$ activation in TSH-differentiated thyroid organoids lead to similar phenotype than cAMP-treated cancer organoids.** Thyroid genes expression levels $^{were}$ obtained when mESC_Nkx2-1/Pax8_bTg_Braf$^{V637E}$_ERT$^2$ cells were differentiated using hrTSH (A). Treatment with 4OHT for 7 days led to downregulation of thyroid maturation genes, *TSHR, Tg, Nis* and *Tpo* when compared to cAMP control organoids (B). Bar graphs represent the median (IQR) of 3 independent experiments with individual values shown. Comparisons were performed against the cAMP condition. (*p<0.05 ; **p<0.01 ; ***p<0.001; ***p<0.0001; Mann-Whitney U test). Nkx2-1 and Tg immunostaining shows that under Braf$^{V637E}$ activation by 4OHT (7 days) thyroid organoids lose the follicular organization while the levels of Tg protein is reduced compared to the cAMP control condition (C). Scale bars, 20 µm.

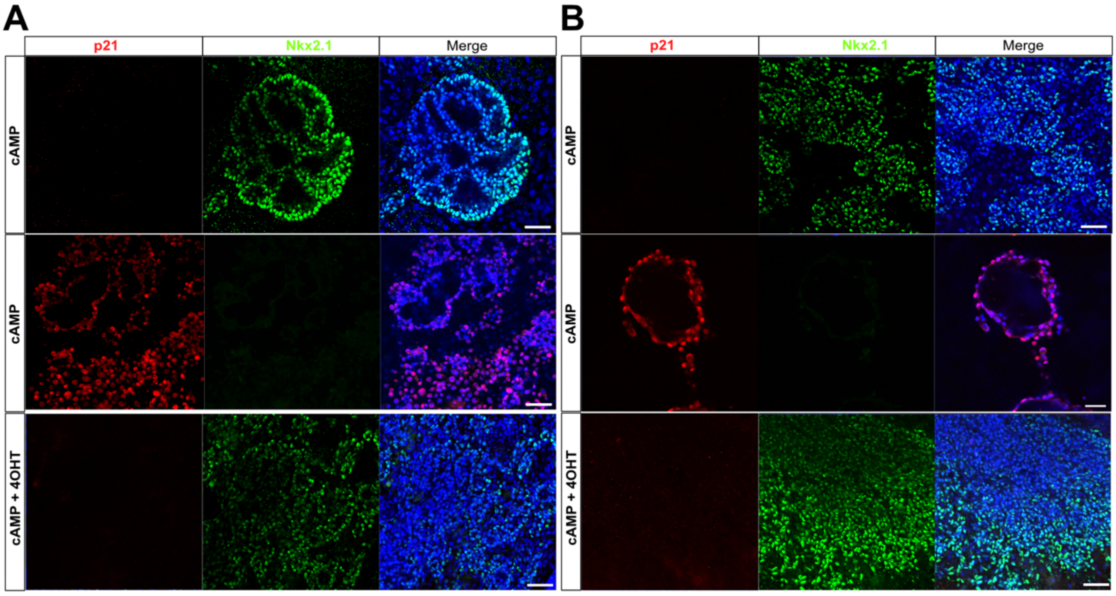


**Supplementary Fig. 5 Braf$^{V637E}$ induction does not induce the expression of p21 in thyroid organoids.** Confocal images show the expression of Nkx2.1 and p21 after 7 (A) and 21 days (B) after induction of *Braf* oncogene by 4OHT treatment. The absence of co-expression of p21 among the Nkx2.1 thyroid cells indicates the absence of oncogene-induced senescence among the cancer organoids. Scale bars, 50 µm.

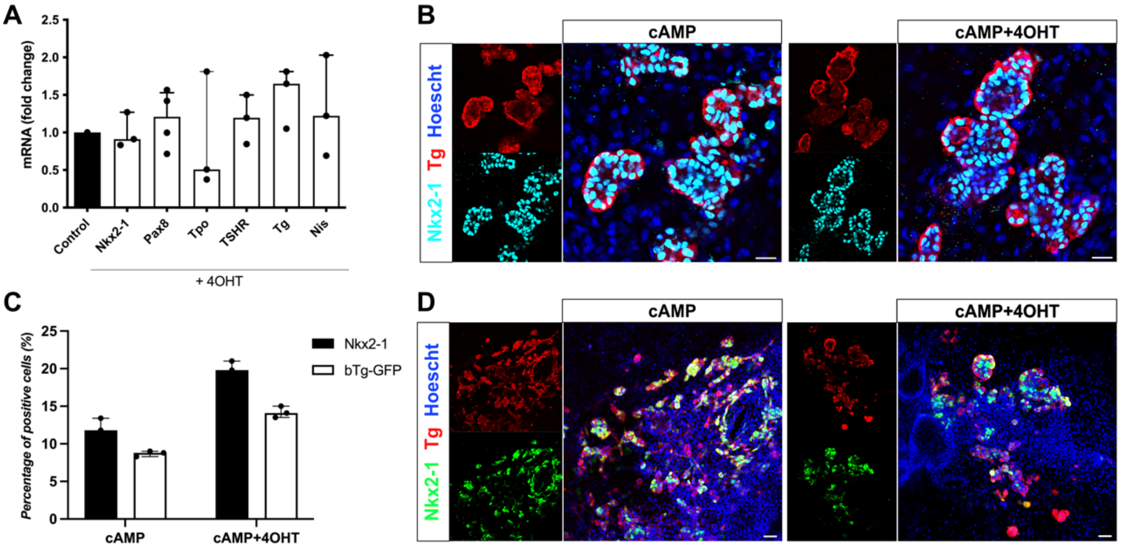


**Supplementary Fig. 6 Treatment with 4OHT in WT thyroid organoids did not lead to cancer phenotype.** TRE-Nkx2-1-Pax8_bTg-eGFP (WT) cell line treated for 7 days with 4OHT did not show changes in gene expression when compared to the control (cAMP) condition (A). Bar graphs represent the median (IQR) of 3 independent experiments with individual values shown. Comparisons were performed against the cAMP (control) condition. (Mann-Whitney U test). Confocal images show that Tg and Nkx2-1 staining and the follicular structures are preserved after 7 days of 4OHT treatment (B). Quantification of the Nkx2-1 and GFP proportions among cAMP and 4OHT-treated (7 days) samples (C). Bar graphs represent the median (IQR) of 3 independent experiments with individual values shown. Comparisons were performed against the cAMP (control) condition. (Mann-Whitney U test). Confocal images show that Tg and GFP co-staining is found only in a proportion of the Tg-expressing cells (D). Scale bars, 50 μm.

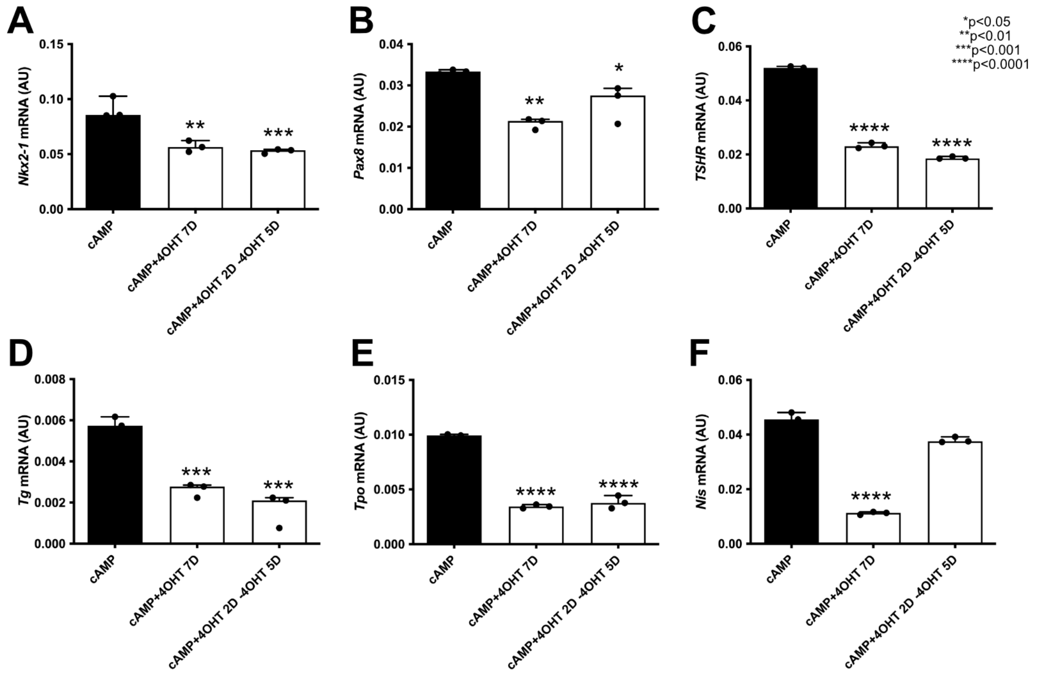


**Supplementary Fig. 7 4OHT interruption does not restore the thyroid gene expression levels.** The induction of Braf$^{V637E}$ for 2 days followed by 4OHT removal for 5 days was not enough to recover *Nkx2-1* (A), *Pax8* (B), *TSHR* (C), *Tg* (D), *Tpo* (E) mRNA to the control levels (cAMP). However, it increased *Nis* expression (F) to control values (cAMP). Bar graphs represent the median (IQR) of 3 independent experiments with individual values shown. Comparisons were performed against the cAMP condition. (*p<0.05 ; **p<0.01 ; ***p<0.001; ***p<0.0001; Mann-Whitney U test).

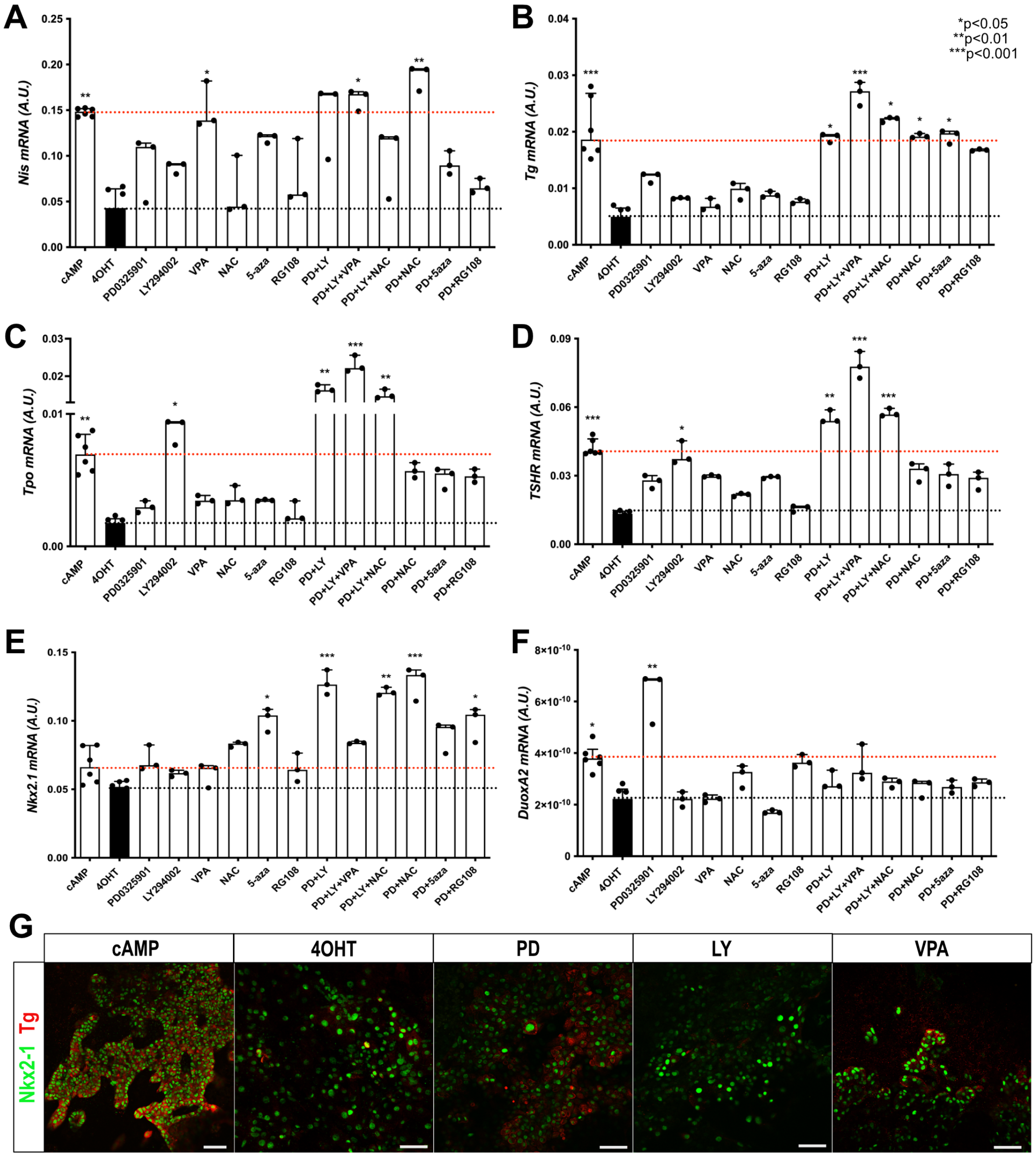


**Supplementary Fig. 8 Effect of distinct classes of inhibitors on thyroid markers.** Gene expression levels of *Nis* (A), *Tg* (B), *Tpo* (C), *TSHR* (D), *Nkx2.1* (E) *and Duox2A* (F) among normal thyroid organoids, Braf$^{V637E}$-expressing (4OHT) and treated with distinct inhibitor (4OHT+inhib). Bar graphs represent the median (IQR) of 3 independent experiments with individual values shown. Comparisons were performed against the 4OHT condition. (*p<0.05 ; **p<0.01 ; ***p<0.001; Mann-Whitney U test). Nkx2-1 and

Tg co-staining shows that isolated PD0325901, LY294002 and VPA treatments are insufficient to revert the inhibitory effect of Braf$^{V637E}$ activation on Tg expression and thyroid follicular organization. Scale bar, 50 µm.

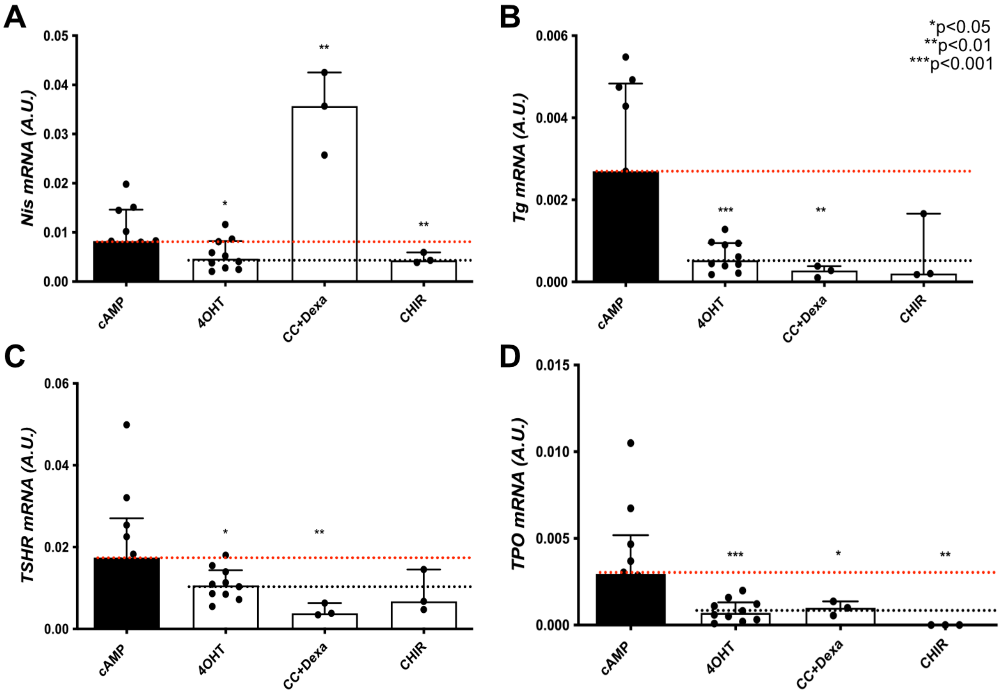


**Supplementary Fig. 9 Inhibition of inflammation and induction of Wnt/β-catenin pathway are not enough to revert thyroid dedifferentiation.** Treatment of cancer organoids (previously treated with 4OHT) with combined anti-inflammatory drugs, dexamethasone, and CC-5013 (TNF-alpha inhibitor) recovered *Nis* mRNA to control (cAMP) levels (A). On the other hand, nor dexamethasone+CC-5013, nor CHIR- 99021 (Wnt/β-catenin activator) could improve the expression levels of *Tg* (B), *TSHR* (C) and *Tpo* (D). Bar graphs represent the median (IQR) of 3 independent experiments with individual values shown. Comparisons were performed against the cAMP condition. (*p<0.05 ; **p<0.01 ; ***p<0.001; Mann-Whitney U test).

### Supplementary tables

**Supplementary Table 1.** Antibodies information.

| Primary antibodies | | | | | | |
|---|---|---|---|---|---|---|
| ***Protein Target*** | ***Provider*** | ***Catalog number*** | ***Host Species*** | ***Dilution Flow Cyt.*** | ***Dilution IF*** | ***Dilution WB*** |
| AFP | Santa Cruz | sc-8108 | Goat | | 1:100 | |
| β-III Tubulin | Eurogentec | MMS-435P-200 | Mouse | | 1:1,000 | |
| αSMA | Abcam | ab32575 | Rabbit | | 1:1,000 | |
| Nkx2-1 | Abcam | ab76013 | Rabbit | 1 :100 | 1:500 | |
| Pax8 | Cell Signaling | 59019 | Rabbit | | 1:500 | |
| Tg | Dako | A0251 | Rabbit | | 1:2,000 | |
| Tg | Abcam | Ab187378 | Mouse | | 1:250 | |
| Nis | Gift from N. Carrasco | | Rabbit | | 1:1,000 | |
| Tg-I | Gift from C. Ris-Stalpers | | Mouse | | 1:250 | |
| E-cadherin | BD Biosciences | 610181 | Mouse | | 1:1,000 | |
| ZO-1 | Invitrogen | 339100 | Mouse | | 1:500 | |
| Hoechst | Invitrogen | 33342 | | | 1:1,000 | |
| p-ERK1/2 | Cell Signaling | 4370 | Rabbit | | | 1:1,000 |
| ERK1/2 | Santa Cruz Biotechnology | sc-514302 | Mouse | | | 1 :400 |
| β-Actin | Cell Signaling | 3700 | Mouse | | | 1:1,000 |
| FITC Rabbit Anti- Active Caspase-3 | BD Biosciences | 559341 | Rabbit | 1 :5 | | |
| APC anti-BrdU | BD Biosciences | 552598 | | 1 :50 | | |
| **Secondary antibodies** | | | | | | |
| Cy3-conjugated | Jackson Immunoresearch | 715-165-150 | Donkey anti-mouse IgG | | 1:500 | |

| | | | | | | |
|---|---|---|---|---|---|---|
| Cy3-conjugated | Jackson Immunoresearch | 711-165-152 | Donkey anti-rabbit IgG | | 1:500 | |
| Cy3-conjugated | Jackson Immunoresearch | 705-165-147 | Donkey anti-goat IgG | | 1:500 | |
| Alexa fluor 488-conjugated | Jackson Immunoresearch | 715-545-150 | Donkey anti-mouse IgG | | 1:500 | |
| Alexa fluor 647-conjugated | Jackson Immunoresearch | 715-605-150 | Donkey anti-mouse IgG | | 1:500 | |
| Alexa fluor 647-conjugated | Jackson Immunoresearch | 711-605-152 | Donkey anti-rabbit IgG | 1:300 | 1:500 | |
| Anti-rabbit IgG, HRP-linked | Cell Signaling | 7074 | | | | 1:1,000 |
| Anti-mouse IgG, HRP-linked | Cell Signaling | 7076 | | | | 1:1,000 |

**Supplementary Table 2.** List of primers used for RT-qPCR analysis.

| **Gene** | **Primer Forward** | **Primer Reverse** |
|---|---|---|
| *β2m-Globulin* | GCTTCAGTCGTCAGCATGG | CAGTTCAGTATGTTCGGCTTCC |
| *Nkx2-1 exog.* | GGCGCCATGTCTTGTTCT | ACACCGGCCTTATTCCAAG |
| *Nkx2-1* | GGCGCCATGTCTTGTTCT | GGGCTCAAGCGCATCTCA |
| *Pax8* | CAGCCTGCTGAGTTCTCCAT | CTGTCTCAGGCCAAGTCCTC |
| *Foxe1* | GGCGGCATCTACAAGTTCAT | GGATCTTGAGGAAGCAGTCG |
| *Tg* | GTCCAATGCCAAAATGATGGTC | GAGAGCATCGGTGCTGTTAAT |
| *Nis/Slc5a5* | AGCTGCCAACACTTCCAGAG | GATGAGAGCACCACAAAGCA |
| *Tshr* | GTCTGCCCAATATTTCCAGGATCTA | GCTCTGTCAAGGCATCAGGGT |
| *Tpo* | ACAGTCACAGTTCTCCACGGATG | ATCTCTATTGTTGCACGCCCC |
| *Braf*$^{V637E}$ *exog.* | GCTTGTGCTTCTCCGAAAAC | AAGGCCAGGCTGTTCTTCTT |
| *Dio1* | ATGGCCAGGAACCCCCG | GCCTGCTGCCTTGAATGAAA |
| *Dio2* | TTCTCCAACTGCCTCTTCCTG | CCCATCAGCGGTCTTCTCC |

| | | |
|---|---|---|
| *Duoxa2* | GCCTGGAATCCGTGGCACTC | TCCCCACGAACCAGTCTCCACT |
| *Gal3* | TTGAAGCTGACCACTTCAAGGTT | AGGTTCTTCATCCGATGGTTGT |
| *Fn1* | AATCACAGTAGTTGCGGCAGGAGA | TCTGTCCCAGGCAGGAGATTTGTT |

**Supplementary Table 3.** Gene list used for thyroid differentiation (TDS and eTDS) and ERK scores calculation.

| | **TDS** | **eTDS** | **ERK score** |
|---|---|---|---|
| **Included** | DIO1<br>DIO2<br>DUOX1<br>DUOX2<br>FOXE1<br>GLIS3<br>NKX2-1<br>PAX8<br>SLC26A4<br>SLC5A5<br>SLC5A8<br>TG<br>THRA<br>THRB<br>TPO<br>TSHR | BCL2<br>CDH16<br>CLCNKB<br>DLG2<br>DPP6<br>FAM167A<br>FHL1<br>FLRT1<br>GPM6A<br>GRIN2C<br>HGD<br>KIT<br>LMOD1<br>LRP2<br>MATN2<br>MPPED2<br>PKHD1L1<br>PLA2R1<br>SLC26A7<br>SLC4A4<br>SORBS2<br>STXBP5L<br>TFCP2L1<br>TFF3<br>TPPP<br>WSCD2<br>ZMAT4 | ARID5A<br>B4GALT6<br>BRIX1<br>BYSL<br>CCND1<br>CD3EAP<br>CHSY1<br>DDX21<br>DUSP4<br>DUSP6<br>EGR1<br>ELOVL6<br>ETV1<br>ETV4<br>ETV5<br>FOS<br>FOSL1<br>GEMIN4<br>GTF2A1L<br>GNL3<br>GPR3<br>GTPBP4<br>HMGA2<br>HYDIN<br>IER3<br>KIR3DL2<br>LIF<br>MAFF<br>MAP2K3<br>MYC<br>NOP16<br>PHLDA2<br>PLK3<br>POLR1C<br>POLR3G<br>PPAN<br>PPAT<br>PYCRL<br>RRS1<br>SEMA6A<br>SH2B3<br>SLC1A5<br>SLC4A7<br>SPRED2<br>SPRY2<br>SPRY4<br>TNC<br>TNFRSF12A<br>TSR1<br>WDR3<br>YRDC |
| **Missing** | | LOC286002<br>MT1G | CXCL8 |

**Supplementary Table 4.** List of the DEGs for each gene enrichment classification.

Included as a separate file.
